# Brain Connectivity Changes in Meditators and Novices during Yoga Nidra: A Novel fMRI Study

**DOI:** 10.1101/2023.09.15.557655

**Authors:** Suruchi Fialoke, Vaibhav Tripathi, Sonika Thakral, Anju Dhawan, Vidur Majahan, Rahul Garg

## Abstract

Yoga Nidra practice aims to induce a deeply relaxed state akin to sleep while maintaining heightened awareness. Despite the growing interest in its clinical applications, a comprehensive understanding of the underlying neural correlates of the practice of Yoga Nidra (YN) remains largely unexplored. In this novel fMRI investigation, we aim to discover the differences between wakeful resting states and states attained during YN practice. The study included individuals experienced in meditation and/or yogic practices, referred to as ‘meditators’ (n=30), and novice controls (n=31). The GLM analysis, based on audio instructions, demonstrated activation related to auditory cues without concurrent Default Mode Network (DMN) deactivation. Additionally, meditators exhibited heightened bilateral thalamic activation compared to novices. DMN seed based Functional connectivity (FC) analysis revealed significant reductions in connectivity among meditators during YN as compared to controls. We did not find differences between the two groups during the pre and post resting state scans. Moreover, when DMN-FC was compared between the YN state and resting state, meditators showed distinct decoupling, whereas controls showed increased DMN-FC. Finally, meditators exhibit a remarkable correlation between reduced DMN connectivity during Yoga Nidra and self-reported hours of cumulative meditation and yoga practice. Together, these results suggest a unique neural modulation of the DMN in meditators during Yoga Nidra which results in being restful yet aware, aligned with their subjective experience of the practice. The study deepens our understanding of the neural mechanisms of Yoga Nidra, revealing distinct DMN connectivity decoupling in meditators and its relationship with meditation and yoga experience. These findings have interdisciplinary implications for neuroscience, psychology, and yogic disciplines.

## Introduction

Yoga Nidra practice, a meditative technique originating from the ancient Indian tradition, has garnered global attention for its potential to improve psychological well-being and health^1^. Despite the growing interest in its clinical applications, a comprehensive understanding of the underlying neural correlates of Yoga Nidra remains largely unexplored^2^. Literally translated as “Yogic Sleep practice,” Yoga Nidra practice stands distinct from many conventional meditation practices in its approach and execution. While most forms of meditation require a seated, upright posture, Yoga Nidra practice is typically attempted in *Shavasana*, a supine position resembling the stillness of a corpse. In contrast to the wakeful, self-regulated focus typically associated with meditation, Yoga Nidra practice involves a series of audio-guided instructions that aim to induce a deeply relaxed state, mirroring the serenity experienced during deep sleep. Even so, throughout the practice, the participant retains conscious awareness, attentively following directions that gently and systematically guide their focus to different parts of the body, breathing, or *mantras*^1^. The practitioner remains in a state of light withdrawal of the 5 senses (*pratyahara*)^3^ with four of their senses internalized and only the hearing still connects to the instructions. The exceptional allure of this technique stems not merely from the deep relaxation and mindful awareness it provides, but also as a method to progressively master entering the most profound states of meditation (*samādhi*)^3–5^. In traditional texts, Yoga Nidra is generally described as a specific meditative state in which advanced practitioners maintain deep rest while maintaining awareness both internally and of one’s surroundings^2,6^.

Yoga Nidra as an audio guided practice for a wide range of individuals and conditions was popularized in the 1960s through the seminal book of Swami Satyananda Saraswati.^1^ Following this, numerous intervention studies have highlighted the notable impact of YN practice on physiological, psychological, and cognitive improvements, making it a promising technique for clinical interventions^2,7^. Specific examples include improvement in symptoms of stress, depression, and anxiety^8–12^, migraine^13^, pain^14^, menstrual abnormalities^15,16^, somatoform disorders^17^, diabetes^18^, and hypertension^11,19^. Further, there is an assertion that one hour of Yoga Nidra (YN) practice can potentially yield benefits equivalent to that of 3-4 hours of traditional sleep^1^. Interestingly, YN practice has also shown benefits in improving the outcomes related to insomnia and sleep quality in several clinical studies^20–23^.

YN has shown a unique ability to be tailored to diverse cultural contexts. For example, a variant of YN known as iRest has been specifically designed to treat Post Traumatic Stress Disorder (PTSD) and the sleep quality of U.S. war veterans^24,25^. Custom iRest protocols are applied to several other therapeutic contexts including occupational stress, sexual trauma, and insomnia^26–28^. More recently, YN practice has been popularized as NSDR (Non-sleep Deep Rest) by Dr./Prof. Andrew Huberman, further expanding its reach and acceptance.

Multiple studies that have looked at the EEG signatures of YN. Parker *et. al*. hypothesized that the ‘perfect’ state of YN would be akin to deep non-REM sleep (exclusive delta-wave predominance) with maintained internal and external awareness, finally culminating into *turīya* (a state devoid of phenomenological content)^6^. Non-peer-reviewed reports of pilot studies have shown that accomplished yogis have demonstrated such states during YN^5,29^. Two pioneering PET-EEG studies reported widespread increases in theta activity during YN (compared to a resting state), indicating a shift towards a more relaxed state^30,31^. The first PET-EEG study with highly experienced yoga practitioners (n=9) revealed bilateral hippocampal activation, along with activation in Wernicke’s area, the occipital lobe, and the parietal and frontal lobes during different stages of YN. Subjectively, the participants described the YN state as “reduced conscious control of attention and behavior, relaxation and a loss of will”, distinctly different from a control condition of resting state^30^. The second PET-EEG study involving 8 experienced Yoga teachers, revealed a 65% increase in dopamine release during YN. This increase was associated with decreased activity in the striatum, as well as reduced executive control and attentional engagement^31^.

Interestingly, a clinical EEG study revealed that compared to the resting state pre-YN, there was an increase in delta power (and no change in theta power) in the central area and a decrease in delta power in the prefrontal area during YN. These findings were attributed to local sleep and not regular sleep during the practice of YN^20^. In a separate EEG study, six novices underwent 12 YN sessions, reporting altered states of consciousness according to the Phenomenology of Consciousness Inventory. Intriguingly, despite each 2-hour YN session, no EEG recordings from any subject showed sleep indicators like K-complexes or sleep spindles^32^.

Although the number of neuroimaging studies on Yoga Nidra is limited and heterogeneous, consisting only of two PET studies and no fMRI study to our knowledge, they provide valuable insights into the state of mind during the Yoga Nidra practice. Along with the EEG studies, these studies have revealed changes indicative of sleep-like relaxation, altered consciousness, and modulation of dopamine release. The guided directions and supine position of YN practice make it accessible for neuroimaging and clinical studies, enabling repeated experiments within its practice. Despite its recognition in wellness and theoretical consciousness studies, it has received limited attention in terms of its neural underpinnings. Existing research and theoretical postulations suggest that YN practice might influence specific neural circuits involved in mind-wandering, self-regulation, and attention. However, these interpretations are mostly inferred from broader meditation research^33^, and the unique characteristics of Yoga Nidra - its blend of sleep-like relaxation with conscious awareness - remain underexplored. This gap in knowledge forms the foundation for our study as we seek to illuminate the neural correlates of Yoga Nidra.

In the context of investigating the neural correlates of Yoga Nidra, the Default Mode Network (DMN) assumes specific importance given its involvement in higher cognition such as internal mentation^34^, self-projection^35^, theory of mind^36^, intrinsic processing^37^, and mind wandering^38^, It lies at the further end of the hierarchy of cognitive networks and evolved when the neocortex expanded^39^. The DMN deactivates during externally demanding tasks in an anti-correlated manner with the Dorsal Attention Network (DAN)^40^. DMN is involved in scene construction which activates when the brain projects itself into the past or the future or is engaged in thinking about others^41^. Alpha and beta wave activity is closely linked with DMN activations^42,43^, and could represent an integrated system. The dysfunction of DMN is associated with various mental disorders like Alzheimer’s disease^44^, anxiety, depression, and ADHD^45^. Meditation studies have shown that meditative practices affect the activity in the DMN, especially in the anterior cingulate cortex (ACC)^46^, and medial prefrontal cortex (mPFC)^47^, and are associated with anxiety relief^46^, emotional stability^48^, and reduced mind wandering^49^.

In this study, we used fMRI to explore the neural correlates of Yoga Nidra practice during a 20 minutes practice of YN in a group of meditators (n=30) and a matched group of novices (n=31). We specifically compare the effects of the YN practice with resting states recorded both before and after the practice of Yoga Nidra. Our investigation delves into how Yoga Nidra practice impacts the DMN’s functional connectivity, and how these changes correlate with the total duration of an individual’s meditation practice. More specifically, in this paper, we aim to find out: Does the functional connectivity within the brain, as captured through seed regions of the DMN during Yoga Nidra practice, resemble the patterns seen during sleep or is it similar to a wakeful rest? And are there differences in DMN interconnectivity between regular meditation practitioners and novices during Yoga Nidra practice and resting state sessions?

## Results

We gathered fMRI and subjective questionnaire data from meditation practitioners (n=30) and demographically matched novice Controls (n=31); see Table S1 for the demographics details. We analyzed the audio-guided Yoga Nidra paradigm (in English in the voice of Sri Sri Ravishankar, founder Art of Living) and pre- and post-Yoga Nidra resting state scans. The Yoga Nidra script used in the fMRI paradigm is represented in Figure 1. The paradigm is divided into four periods: T1 to T3 each lasting 240 seconds, and T4 spanning 215 seconds for functional connectivity analysis. The instructions systematically guide the practitioner’s attention across different body parts followed by a mantra chant and period of silence, thereby fostering a relaxed state.

**Figure 1.**
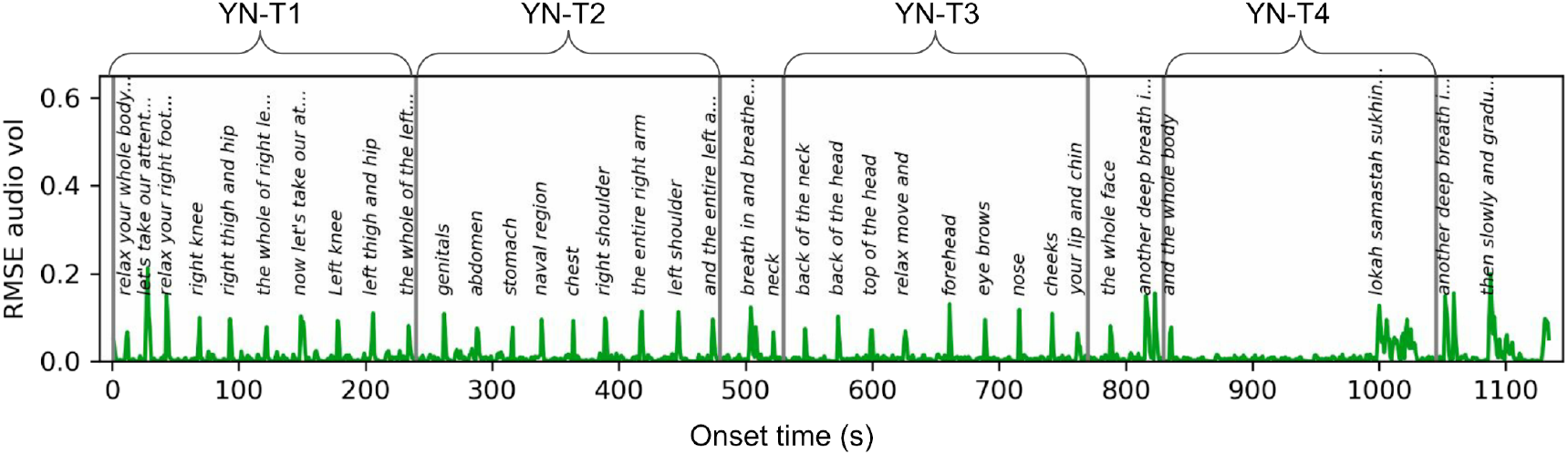
Illustration of the Yoga Nidra audio-guided paradigm. The x-axis represents the progression of subtitles over time, while the y-axis denotes the RMSE volume of the audio instructions. The practitioner’s attention is sequentially guided to different body parts as depicted in the figure. The paradigm is divided into four periods: T1 to T3 each lasting 240 seconds, and T4 spanning 215 seconds for connectivity analysis chosen to exclude any explicit breathing instructions.

### Phenomenological Self-Reports of Yoga Nidra Practice

Both groups were instructed to practice Yoga Nidra (same script as the one used during fMRI) 2-3 times in the weeks prior the scanning day. Most meditators (16/30) and controls (22/31) indicated practicing 1-5 times in the past month with two controls and five meditators reported practicing 6-10 times and 3 meditators reported practicing >10 times in the past month. Seven controls and five meditators reported no practice in the past month and remaining didnt respond. We also collected the subjects’ subjective experience during the scan, including the number of times they think they fell asleep, and restfulness. The survey results indicated, most control participants (21/31) reported falling asleep < 2 times, with a smaller number (4/31) falling asleep 3-4 times, and 4 participants experienced constant falling in and out of sleep. Similarly, in the meditators group, a majority (25/30) reported falling asleep < 2 times, one fell asleep 3-4 times, two reported constantly falling in and out of sleep, and one reported sleeping mostly. The remaining subjects did not respond to this question. Regarding the experience of Yoga Nidra in the scanner, in the control group, 17 participants reported being uncomfortable during the practice, while 3 reported feeling deeply relaxed, and 7 felt rested. Comparatively, among meditators, only 3 reported discomfort during the practice, 8 felt deeply relaxed, and a majority (18) reported feeling rested. The data suggests that the meditators found the Yoga Nidra experience in the scanner more comfortable and restful as compared to the control group.

### BOLD Activations Modeled after Yoga Nidra Auditory Instructions

We performed the General Linear Model (GLM) analysis with the aim to model the BOLD response in relation to the audio instructions. Accordingly, the explanatory variable (EV) in the model was the presence of audio instructions, coded as 1 (refer to Fig. 1 for detailed timing of the instructions). Group-level activations for meditators and controls (n=61) observed during the practice of Yoga Nidra based on General Linear Model (GLM) analysis are presented in Figure 2. Additionally, in Table S2, we provide a list of significant clusters of activations and deactivations. YN practice initiates activations in multiple brain regions specifically, the temporal gyri, responsible for auditory and language processing, are activated, highlighting the key role of auditory reception and comprehension during YN practice. The supplementary motor area (SMA) and the postcentral gyrus (PCG) regions that are involved in motor planning and control are activated as expected as the instructions involve scanning of body parts. Other areas include the thalamus and brainstem, crucial for the relay of sensory and motor signals^50^. Activations in the anterior cingulate cortex (ACC) well-known for its role in both cognition and emotion is notable.^51^ Similarly, activation in the insula, inferior frontal regions, cerebellum, and brainstem, key regions involved in emotional processing and cognitive control were observed. Lastly, superior frontal gyrus, contributing to high-level cognitive functions, also exhibit activation. However, deactivation in several regions, including the frontal pole, superior frontal gyrus, and other areas may suggest a decreased demand for some complex cognitive processes, reinforcing the relaxing and introspective nature of YN.

**Figure 2.**
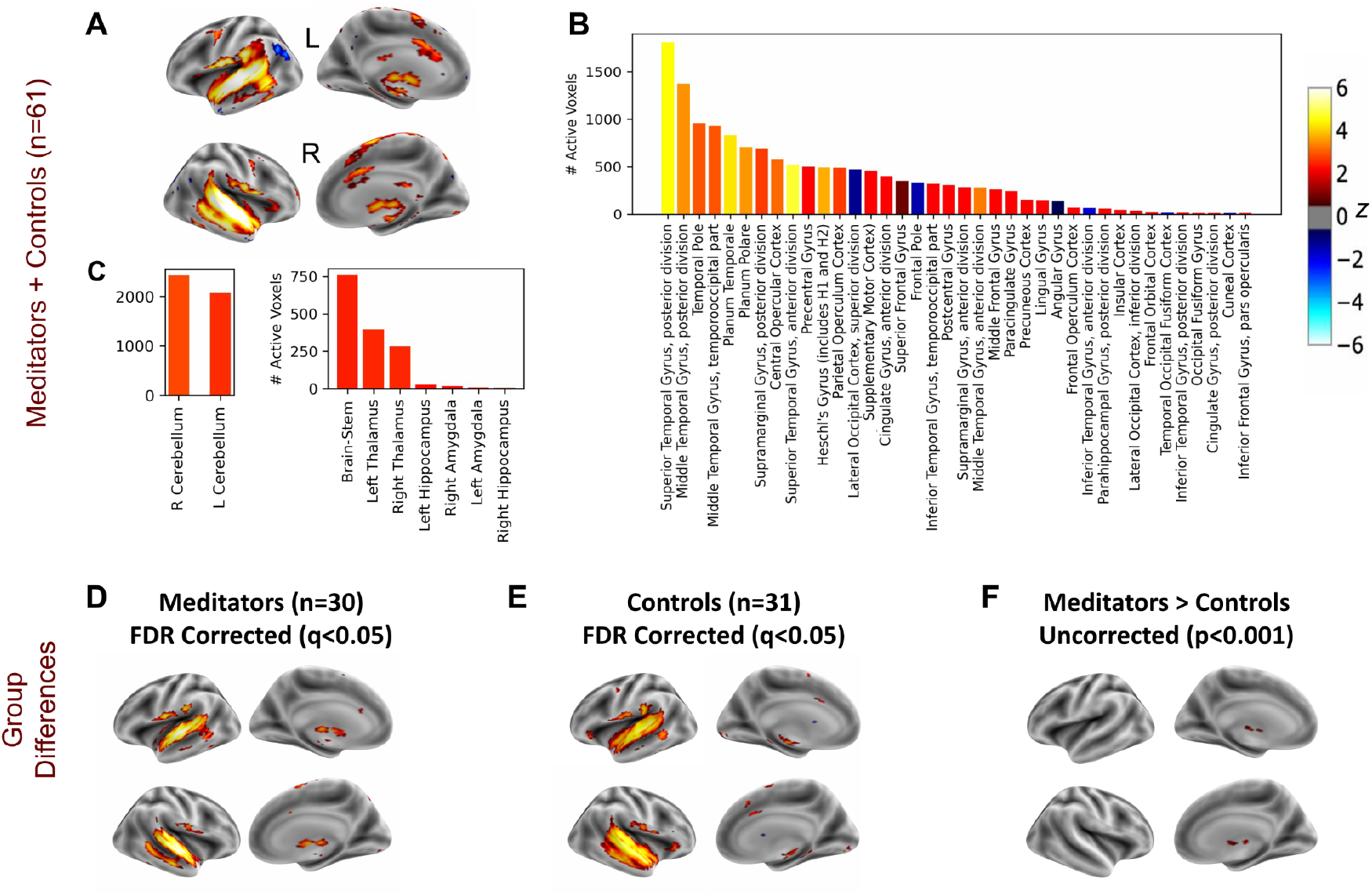
Brain activations during Yoga Nidra, show stimulated auditory, language, motor, and limbic regions, and distinctive thalamic activation in meditators. (A) Surface visualizations of z-statistics, adjusted using False Discovery Rate (FDR) correction (q < 0.05), derived from GLM analysis of Yoga Nidra, all subjects (meditators and controls collectively). The explanatory variable (EV) was modeled with the presence of any audio instruction as 1 (Fig. 1). (B) Active voxel counts for Cortical ROIs from the above map are presented (the color of the bars represents the average z-score within that ROI). Consistent with the auditory instructions involved in Yoga Nidra (YN), which guide attention to different body parts, activations were observed not only in the auditory and language regions but also in motor, supplementary motor, and somatosensory areas. Regions associated with memory and emotional processing are also active including Insula and Parahippocampal Gyrus. Interestingly, despite activation of language and auditory areas, DMN regions do not exhibit concurrent de-activation. (C) Active voxel counts for Cerebellum and Subcortical ROIs exhibiting activations in limbic regions including the brainstem, Thalamus, and Amygdala (the color of the bars represents the average z-score within that ROI). (D) Surface plots of z-stats, FDR-corrected (q < 0.05), from GLM analysis of Yoga Nidra for meditators (n=30) and (E) same for control subjects (n=31). (F) Although the FDR-corrected maps (q < 0.05) from the comparison ‘Meditators>Controls’ didn’t have any surviving voxel, uncorrected maps with a threshold at p<0.001 underscore bilateral activation in the thalamic regions.

Figures 2(D-F) offer a comparative view of the BOLD response to YN instructions in meditators versus controls. The intriguing observation of thalamic activation in meditators, absent in the controls is notable. Remarkably, despite the activation of language, and auditory areas, as participants attended to the instructions, DMN regions did not exhibit concurrent de-activation. It could either be due to a sustained level of engagement during the YN practice, preventing the deactivation of DMN, or due to a reduction in mind wandering/autobiographical processes during the YN practice, or due to the lack of perfect synchronization of DMN deactivation with auditory YN instructions^52^. While this could potentially contribute to its deep relaxation effect, associating the lack of DMN deactivation directly with increased relaxation would require further investigation.

### DMN Connectivity During Yoga Nidra

#### Group DMN FC During Yoga Nidra and Resting States

For the seed based functional connectivity analysis, we selected the following seeds within the Default Mode Network (DMN): the Posterior Cingulate Cortex (PCC), medial Prefrontal Cortex (mPFC), Right Inferior Parietal Lobule (right-IPL), and Left Inferior Parietal Lobule (left-IPL)^53^. We extracted temporal sequences from a 10 mm-radius sphere centered on these seeds, and computed pairwise correlations with all brain voxels, followed by Fisher’s-Z transformation for group-level analysis. The outcomes are displayed on surface maps in Figure S1-S4 for all four DMN seeds in both meditators and controls. The presence of a functional DMN manifested during both resting states and Yoga Nidra (YN). Notably, discernable patterns of functional connectivity (FC) emerged between meditators and controls during the Yoga Nidra practice for each DMN seed. This divergence prompted an exploration of contrasts between the two groups and the stages of Yoga Nidra practice versus resting states discussed in the subsequent sections.

#### Comparison of the DMN FC in Meditators and Novices

Figure 3 suggests differences in the brain activity patterns between meditators and non-meditators during Yoga Nidra (YN). Specifically, during all four stages of YN, meditators exhibit markedly reduced functional connectivity (F(5,341) = 4.84, p < 0.0001, see table S7 for the detailed complete ANOVA table) within the four DMN ROIs, PCC (Figure 3A, Table S3), mPFC (Figure 3B, Table S4), right-IPL (Figure S5A, Table S5), and left-IPL (Figure S5B, Table S6). Interestingly, this pattern of reduced DMN connectivity emerges in the first phase, T1, itself, which lasts only four minutes suggesting that the reduced functional connectivity within the DMN seen in meditators is indicative of a state of deep relaxation and not due to sleep. Moreover, no significant post-hoc differences (p = 0.886) were detected in the functional connectivity of these DMN components between meditators and non-meditators during resting states before or after YN (see table S3-S6 for pairwise connectivity differences). This suggests that these changes in DMN connectivity are not simply inherent characteristics of the meditators’ brains but appear to be specifically tied to the practice of YN. This finding emphasizes the potential of practices like YN to influence brain function in a context-dependent manner, contributing to our understanding of the neural mechanisms that underlie the benefits of meditation and other mind-body practices.

**Figure 3.**
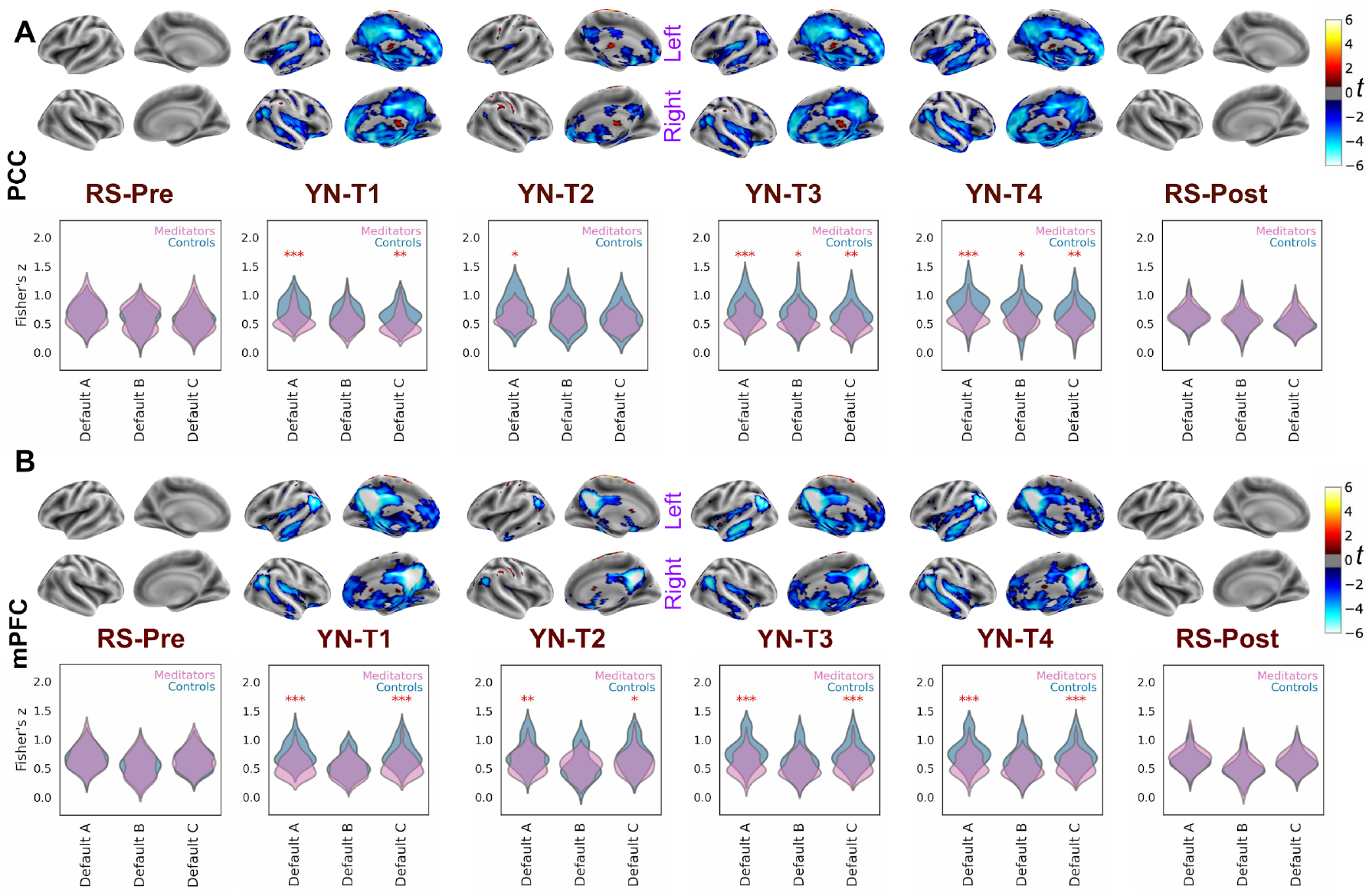
Group Differences in DMN-FC between meditators and controls during Yoga Nidra but not during resting states. The figure uses two-sided t-tests to compare FC in meditators (n=30) and controls (n=31) using DMN seeds (A) Posterior Cingulate Cortex (PCC) and (B) medial Prefrontal Cortex (mPFC). The surface plots display the t-values, corrected for multiple comparisons (FDR-corrected, q < 0.05), across the four stages of Yoga Nidra (YN), as well as in resting states pre and post-YN. The accompanying violin plots illustrate the distribution of Fisher’s z-values for both groups during these stages. These plots represent the averaged Fisher’s z-values of the seeds within three distinct Default Mode Network (DMN) subdivisions (Default A, Default B, and Default C), as outlined by the Schaefer Cortical Atlas. The width of the violin plot at any given y-value (Fisher’s z-value) represents the proportion of data located there, providing a visual representation of the data’s distribution (* represents statistical significance with p < 0.05, ** with p < 0.01 and *** with p < 0.001). Notably, meditators demonstrate significantly reduced DMN connectivity for both PCC and mPFC seeds during all four stages of Yoga Nidra, indicating a deeper state of focused relaxation in experienced meditators. Conversely, no significant differences were observed in the functional connectivity of these seeds in either of the resting states.

#### DMN during Yoga Nidra versus during Resting State

In the context of both meditators and controls, we contrasted the FC during stages of YN with the FC during the resting state prior to YN (Pre-YN). For PCC (Figure 4A), mPFC (Figure 4B), right-IPL (Figure S6A), and left-IPL (Fig S6B) seeds—core components of the DMN—the findings are strikingly divergent between the two groups. Meditators exhibit a marked DMN FC reduction (p < 0.05) throughout Yoga Nidra (T1 to T4) compared to their resting state, potentially signifying YN-induced tranquility with minimized mentation and autobiographical memory. Contrarily, control subjects display a significant increase (p = 0.01) in DMN FC during T4 compared to the resting state. In the SI (Figure S7), we also include the intra-DMN FC by looking at the correlation between the DMN seed pairs. It is interesting to note that the intra DMN FC is significantly reduced (p < 0.05) during YN compared to the resting state for meditators but not for controls. However, for both meditators and controls, the resting state post YN, wasn’t significantly different (p = 0.5) from the resting state prior to YN, indicating that the changes are context dependent. These findings further underscore the potential of regular meditation and yoga training to modulate brain networks and cognitive states during relaxation.

**Figure 4.**
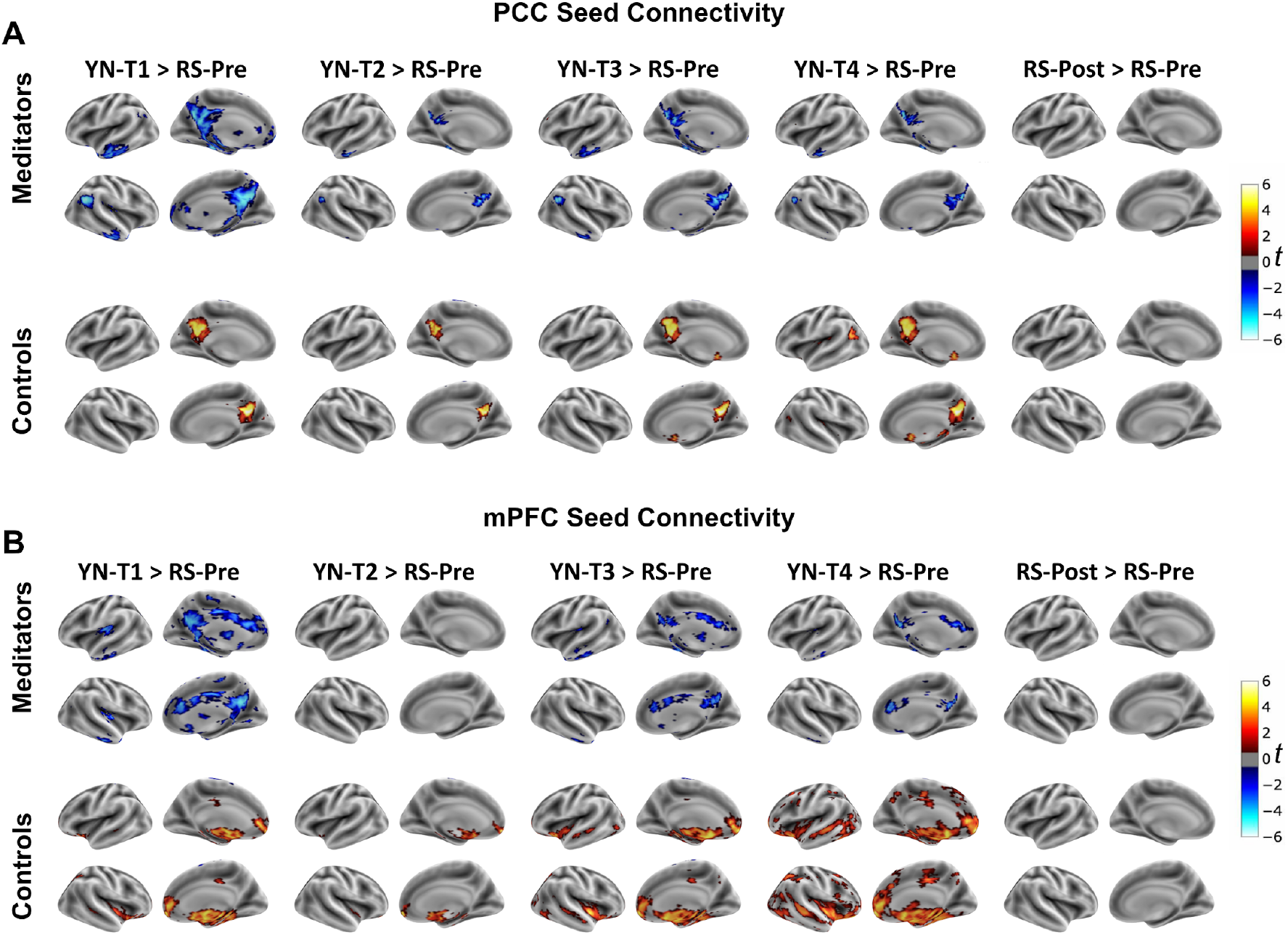
Different functional connectivity changes emerge in meditators and controls during Yoga Nidra compared to rest. For (A) PCC and (B) mPFC seeds, a comparison of the FC during YN (T1 through T4) and the resting state post-completion of Yoga Nidra (RS-Post) is performed with resting state pre-Yoga Nidra (RS-Pre) as a baseline. The surface maps present t-values from two-sided t-tests, corrected for multiple comparisons (FDR-corrected, q < 0.05), providing a visual representation of FC differences. The figure highlights that meditators demonstrate a significant decrease in DMN connectivity during YN compared to their resting state. Conversely, controls display a significant increase in connectivity during YN relative to the resting state.

#### Relationship between total meditation practice and DMN-FC

Figure 5A highlights a compelling negative correlation between the total duration of meditation practice and the functional connectivity (FC) of the medial prefrontal cortex (mPFC) seed with other regions of the DMN (Default A as defined by the Schaefer Atlas^54^) during the Yoga Nidra (YN) practice (YN-T1: r(29) = -0.43, p = 0.006; YN-T2: r(29) = -0.4, p = 0.012; YN-T3: r(29) = -0.41, p = 0.008; YN-T4: r(29) =-0.35, p = 0.025). This intriguing pattern suggests that as meditation experience increases, there is a corresponding decrease in functional connectivity between the mPFC seed and DMN regions during YN. This could potentially signify a trait-like effect, where self-referential thought processes become less dominant during the practice of YN. Interestingly, this linear relationship is absent during the resting states before and after YN, emphasizing the context-dependent nature of this correlation.

**Figure 5.**
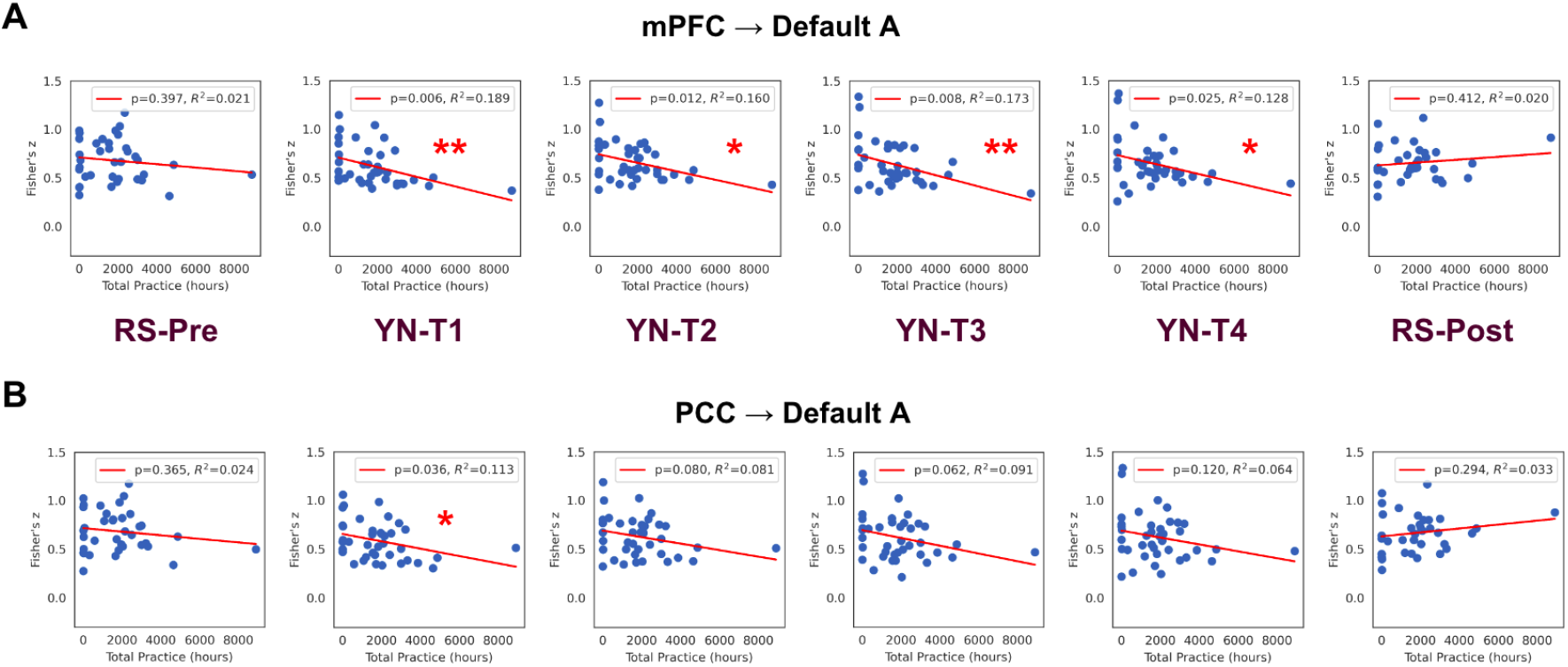
Increased meditation experience significantly correlates with reduced functional connectivity between the mPFC seed and DMN regions. (A) A significant negative correlation between the total duration of meditation practice and the functional connectivity (FC) of the medial prefrontal cortex (mPFC) with other regions of the Default Mode Network (DMN, Default A as defined by the Schaefer Atlas) during the Yoga Nidra (YN) practice. * represents statistical significance with p < 0.05 and ** with p < 0.01. This pattern suggests that increased meditation experience correlates with reduced functional connectivity between the mPFC seed and DMN regions during YN, potentially reflecting a trait-like effect of diminished self-referential thought processes during this meditative state. Interestingly, no such linear relationship was detected during the resting states before and after YN, emphasizing the context-dependent nature of this association. (B) For the Posterior Cingulate Cortex (PCC), similar qualitative trends were observed, albeit with less statistical significance, indicating a potential commonality in the effects of prolonged meditation practice on the functional connectivity of core DMN regions.

To ensure the robustness of our findings, we repeated the analysis after removing the outlier subject with more than 8000 hours of cumulative practice (Figure S8). Remarkably, the results remained significant for all four stages of Yoga Nidra (YN-T1: r(29) = -0.41, p = 0.011; YN-T2: r(29) = -0.36, p = 0.024; YN-T3: r(29) = -0.36, p = 0.031; YN-T4: r(29) = -0.35, p = 0.031), reinforcing the strength and reliability of our initial findings. Deploying the PCC Seed (Figure 5B), and bilateral IPL seeds (Figure S9) also reveals similar trends, although their corresponding p-value is greater. This observation suggests a potential shared influence of prolonged meditation practice on the functional connectivity of core DMN seeds.

## Discussion

Yoga Nidra, a unique practice combining detailed body scanning with deep relaxation and heightened awareness, is subjected to fMRI scrutiny for the first time in our research. This investigation involves both regular meditators and controls, aiming to uncover the specific neural activations and connections at play during Yoga Nidra. Central to our inquiry is determining its distinction from wakeful rest states and comparison between meditators and controls. Resting state scans conducted before and after Yoga Nidra sessions provide suitable control conditions.

Simultaneously, the wealth of existing research functional connectivity of DMN during various sleep stages offers a comparative landscape.

GLM analysis modeled after presence of audio instruction, revealed activations in regions involved in auditory and language comprehension, as well as motor planning in both groups. Furthermore, we note activations in areas associated with emotional regulation including the ACC, insula, and the limbic system. This observation aligns with the Swami Satyananda Saraswati’s proposition that “Yoga nidra develops control of the emotional reactions and autonomic responses”^1^.

Our analysis additionally uncovered an intriguing distinction between meditators and controls - meditators exhibit pronounced bilateral thalamic activation, a response absent in controls. This finding gains significance when juxtaposed with research that delved into the neural response to auditory stimuli during various stages of sleep employing fMRI and EEG recordings^55^. The study unveiled bilateral activation in the auditory cortex, caudate, and thalamus, during wakefulness and nonrapid eye movement (NREM) sleep. However, they noted a marked reduction in the activation of the thalami during NREM sleep compared to wakefulness^55^. The thalamus, often considered the ‘gateway to consciousness’, plays a pivotal role in relaying sensory information and maintaining arousal^50,56^. Therefore, its increased activation in meditators may reflect a heightened state of awareness, even amidst the deep relaxation characteristic of Yoga Nidra.

The GLM analysis, however, revealed no deactivation in DMN regions, despite concurrent activation in areas associated with the auditory instructions (Figure 2). There could be two explanations for this. First, Yoga Nidra practice could inherently differ from typical attention-demanding tasks, i.e. the guided and calming nature of Yoga Nidra might foster a degree of engagement that supports the maintenance of DMN connectivity. The absence of DMN deactivation during Yoga Nidra in our study suggests a unique state of consciousness distinct from traditional focused attention (FA) meditation^33,57^. Alternatively, it is possible that disengagement of the DMN regions following the audio instruction might not be strictly stimulus-locked and could exhibit significant variability both across instructions and between subjects. A recent study reported diminished stimulus-locked synchronization in DMN regions in a variety of tasks^52^.

In meditators, significantly reduced FC was observed for each DMN seed (PCC, mPFC, right-IPL, left-IPL) compared to novices at every stage of Yoga Nidra. The body of research comparing the DMN FC of meditators and control groups during meditation is sparse, and notably, no studies have specifically investigated Yoga Nidra. Particularly, Brewer et. al. revealed that compared to controls, expert meditators showed enhanced connectivity between the PCC seed and ‘task-positive’ areas (dACC and dlPFC) across three different meditation types as well as during resting state. This increased connectivity with the dACC and dlPFC was not seen using the mPFC as the seed^58^. The difference between meditators and novices during YN may be due to the unique nature of Yoga Nidra which uses guided instructions to evoke a deep relaxation akin to profound sleep, unlike traditional meditation’s wakeful nature. In a seminal review and meta-analysis of fMRI meditation studies, Fox. et. al.^33^ analyzed common styles of meditation as focused attention, mantra recitation, open monitoring, and compassion/loving-kindness and suggested differences for three others (visualization, sense-withdrawal, and non-dual awareness practices) while keeping Yoga Nidra in a different but unexplored category^33^.

Interestingly, no significant difference between meditators and novices was noted in the resting states either pre or post-Yoga Nidra. Several studies on the resting state have shown that prolonged and regular meditation practitioners have altered DMN FC compared to novices^58–60^. Whereas, some have found no significant differences in DMN FC between meditators and controls during rest^61^. We conjecture that the alteration in traits may necessitate a more extended period of practice and could also be influenced by the demographics and lifestyle of the study population and the specific type of meditation practiced. Notably, yoga practitioners involved in our research are regular householders with an average of 2,827 ± 2,000 hours of yoga practice (which includes yoga postures, breathing, meditation), whereas previous studies documenting changes in resting state FC predominantly focus on meditators (usually renunciate monks) with an excess of 10,000 hours of meditation practice.

When comparing any stage of YN with the resting state, meditators demonstrate a decoupling in DMN connectivity with several brain regions, whereas controls display an increased coupling of the DMN. These divergent patterns potentially reflect the ability of experienced meditators to regulate their mind and brain states during deep stages of relaxation, in contrast to non-meditators who perhaps experienced boredom, discomfort, or increased mind-wandering. In this context, we refer to simultaneous EEG-fMRI studies which have investigated sleep depth effects. As per these studies, sleep depth— gauged through EEG features such as theta activity (4–8Hz) or slow-wave activity (SWA) (< 4.5Hz)— reveals sustained DMN functional connectivity during light sleep, but this connectivity, inclusive of DMN, disintegrates during deep sleep^62,63^. Nevertheless, given the evident reduction in FC of meditators during the initial phase T1, attaining a veritable state of deep sleep appears improbable. This conclusion finds reinforcement in the phenomenological accounts of the meditators, with the majority reporting not experiencing sleep during the practice. Most also meditators reported feeling “deeply relaxed” or “rested” during the practice. This observed decline in FC of meditators underscores the potential of YN practice in modulating the DMN, shifting the brain from a potentially distractible, mind-wandering state to an awake yet deep-sleep like relaxed state.

Remarkably, a significant correlation is observed between the total duration of meditation practice and a reduction in functional connectivity of DMN seeds with DMN regions during Yoga Nidra. As illustrated in Figure 5, we present scatter plots that vividly depict this relationship, with DMN seeds’ functional connectivity, averaged within DMN regions, plotted against the accumulated hours of meditation practice. Notably, these alterations appear to predominantly impact the connectivity between the mPFC seed and DMN regions. Given that the mPFC is associated with self-referential thought processes, this reduced connectivity might be indicative of reduced self-related thinking or mind-wandering during YN among experienced meditators. For the PCC and the bilateral IPL seeds, a similar trend was noted but with less statistical significance. This relationship, again, is not observed during resting states, highlighting the context-dependent nature of these meditation-related changes in brain function. Together, these results seem to provide evidence of a trait-like effect of meditation. As one’s experience in meditative practices grows, we observe discernible changes in the functional connectivity of key DMN regions during the practice of meditation. Such insights highlight the transformative power of regular meditation practices, especially when combined with techniques like Yoga Nidra.

This study opens up thrilling new avenues for future research, for instance, in exploring pathways associated with Yoga Nidra’s efficacy in contexts of neuropsychiatric disorders linked to hyperconnected DMN function^64,65^. When dealing with clinical populations, it’s common to discover that various meditation / yoga techniques have varying degrees of effectiveness for specific disorders and hence, effectively choosing the most suitable practice for an individual patient is a crucial step^66^. The results of this study can provide valuable insights for guiding mental health interventions and enhancing our understanding of the mechanisms underpinning ongoing clinical investigations. Several clinical studies already indicate that the Yoga Nidra practice may mitigate anxiety, stress and depression^9–11,67^. and many have shown promising results of iRest for the same^26,28,68,69^. Also, the promising results of iRest in PTSD, demonstrated by many studies^27,70^, offer compelling evidence to support the belief that Yoga Nidra should prove beneficial for the same. Further research into the mechanisms underlying Yoga Nidra’s impact on neuropsychiatric disorders, mental health and well-being is warranted and holds significant promise.

### Limitations and Future Directions

One limitation of our study is the standalone use of fMRI, which, while providing detailed spatial information, lacks the temporal resolution of EEG. This may have affected our ability to capture rapid changes in neuronal firing and synchrony during Yoga Nidra. Future studies could enhance our understanding of Yoga Nidra by incorporating simultaneous EEG and fMRI recordings. This would allow for a more detailed examination of the spatiotemporal dynamics across various stages of Yoga Nidra and their distinction from different phases of sleep. Furthermore, incorporating a longitudinal study design is critical to better understand the temporal effects and long-term implications of Yoga Nidra on brain activity. This design will enable us to monitor changes over time and elucidate whether regular practice induces any structural or functional brain changes, providing a deeper understanding of the enduring impacts of Yoga Nidra. It is worth noting that both the meditators and controls groups in our study had only 16% females, which could be addressed in future research to ensure gender diversity.

## Conclusion

In this study, we underscore several important insights into the neural underpinnings of Yoga Nidra, a unique yogic practice. Firstly, our GLM analysis, which was modeled using the presence of audio instructions, revealed no deactivation in DMN regions, despite concurrent activation in areas associated with the auditory instructions. Secondly, our findings highlight a significant reduction in DMN functional connectivity among meditators compared to novices across all stages of Yoga Nidra. Parallelly, the GLM modeling of auditory stimuli produced higher activation in bilateral thalamus for meditators compared to novices. This suggests a unique neural adaptation in meditators during Yoga Nidra which results in being restful yet aware. Interestingly, no such difference in DMN functional connectivity was observed between meditators and novices during resting states, either pre or post-Yoga Nidra. Further, when juxtaposing any stage of Yoga Nidra with the resting state, we observed a decoupling in DMN connectivity among meditators, in stark contrast to controls who displayed an increase. This divergence potentially mirrors the advanced ability of meditators to regulate their mind and brain states during deep stages of relaxation, setting them apart from non-meditators. Finally, our analysis reveals a crucial finding: a significant correlation between the total duration of meditation practice and a reduction in functional connectivity of DMN seeds to DMN regions during Yoga Nidra. This suggests that sustained yoga practice may lead to enduring changes in the functional connectivity of DMN. Our findings set the stage for future research to delve deeper into the cognitive and perceptual correlates of Yoga Nidra, particularly how its practice influences DMN connectivity in a context-dependent manner. Moreover, this research holds significant interdisciplinary implications, bridging the gap between neuroscience, psychology, and the study of yogic practices. The insights gained from this study could potentially inform therapeutic interventions in mental health, providing a scientific basis for the integration of Yoga Nidra in stress management and cognitive therapy programs.

## Materials and Methods

### Participant Recruitment and Experimental Design

The study engaged local meditation practitioners in Delhi’s National Capital Region (NCR), utilizing community contacts for recruitment. Participants were classified into two groups: Meditators with atleast 1 year of regular yoga or meditation experience, and Controls, novice-meditators or non-meditators, age and gender-matched with Meditators, with less than 6 months of yoga or meditation experience (see Table S1 for Demographics). In this context, the term “meditators” pertains to subjects with at least 1 year of regular practice that included *asana* (yoga postures), *pranayama* (breathing practices), and meditation. Criteria for inclusion encompassed ages 18-60, a minimum of 10th standard education, English proficiency, and willingness to provide informed consent. Exclusion criteria encompassed known psychiatric illnesses (as assessed by Mini International Neuropsychiatric Interview (MINI) screen), chronic mental conditions, and unwillingness to sign the informed consent form.

The overall study paradigm included two guided meditations: a 20-minute Yoga Nidra protocol and a 23-minute Body Scan protocol, both forms of progressive muscle relaxation methods. Each meditation had a 5-minute eyes-closed resting state run before and after to track functional connectivity changes due to meditation. We also included a paced breathing run in between which allowed the subjects to come from a meditative state to awake state. For this paper, we focussed on the analysis of the Yoga Nidra (YN) practice. The order of the practice was as follows: Structural scan – Resting State (RS) – Yoga Nidra / Body Scan – RS – Rhythmic Breathing – RS – Yoga Nidra / Body Scan – RS. The Yoga Nidra and Body Scan orders were counterbalanced across subjects to negate order effects.

Participants followed the audio-guided Yoga Nidra practice in an eyes-closed supine position for 20 minutes, the transcript is provided in Fig.1 and the audio (in English in the voice of Sri Sri Ravishankar, founder Art of Living) is publicly available [here](https://www.youtube.com/watch?v=zLJu3wQA1Ko&ab_channel=GurudevSriSriRaviShankar).

### Ethical Consideration

The study was reviewed and approved by the Institute Ethics Committee (IEC) of Indian Institute of Technology, Delhi on 22/09/2017 (Ethics application: P-014). The participants provided their written informed consent to participate in this study.

### Data availability

The data of all subjects will be avialble with a subsequent data publication.

### Data Acquisition

We acquired our data on a Philips Ingenia 3.0T whole-body scanner at Mahajan Imaging, Gurugram. We used a T1-weighted 3D MP-RAGE protocol (FOV = 250 mm x 250 mm, flip angle = 8°, acquisition voxel size = 1 mm × 1 mm, slice thickness = 1 mm, 250 slices, slice gap=-0.40 mm) to collect high-resolution anatomical data. The Blood oxygen level-dependent (BOLD) functional data during YN paradigm was acquired using the multi-band echo-planar imaging pulse sequence (TR = 1000 ms, TE=25 ms, flip angle= 65°, FOV = 219 mm x 219 mm, acquisition voxel size = 3 mm x 3 mm, slice thickness = 4 mm, 48 slices without slice gap). The initial 26 subjects were acquired with a multi-band factor of eight and the remaining with the factor three (exact factors per subject listed in Table S8). The functional data during Resting states for all subjects had TR = 1000 ms, TE=25 ms, 65° flip angle, FOV = 219 mm x 219 mm, voxel size = 3×3×4 mm, 60 slices without slice gap, and a multi-band factor of 4.

### Preprocessing

High-resolution T1-weighted were preprocessed using FSL’s BET (frac=0.3) to remove non-brain tissues. Subsequent preprocessing of functional MRI (fMRI) data involved FSL’s MCFLIRT for motion correction and temporal high-pass filtering (cut-off: 100s). Images were then registered using FSL’s FLIRT with subject-specific anatomical images (7-DOF) and the MNI152 standard space (12-DOF)^71^. The slice-time correction was performed to compensate for differences in acquisition timing across slices for each volume.

### Anatomical Segmentation and Noise Regression

The FMRIB Automated Segmentation Tool (FAST) from FSL was employed to extract the cerebrospinal fluid (CSF) tissues from the T1-weighted images^71^. The generated CSF mask was then intersected with a standard ventricle ROI. A noise vector (csf_ts) was calculated from the average signal of the CSF x ventricle mask. Head motion parameters—three rotational (x, y, z in radians) and three translational (x, y, z in millimeters) for each volume—were derived from the MCFLIRT motion correction outputs. These six head-motion parameters, along with the CSF signal, constituted seven noise vectors that spanned the noise subspace. The component orthogonal to the noise vectors (i.e., perpendicular to the noise basis) was employed for subsequent analyses.

### Spatial Smoothing

Preprocessed and de-noised fMRI data were subjected to spatial smoothing using a Gaussian kernel with a 5 mm full-width at half-maximum (FWHM) to enhance signal-to-noise ratio and accommodate intersubject variability.

### General Linear Model

The General Linear Model (GLM) analysis was applied to the pre-processed fMRI data using FSL’s FEAT (FMRI Expert Analysis Tool). The explanatory variable (EV) in the model was the presence of audio instructions, coded as 1 (refer to Fig. 1 for detailed timing of the instructions), with the aim of modeling the BOLD response in relation to the audio instruction. The model was convolved with the canonical hemodynamic response function (HRF), and temporal filtering was applied to match the filtering performed during preprocessing. Following the first-level analysis performed for each subject, the resulting contrasts of parameter estimates (COPEs) were used to obtain group-level statistics. Employing Nilearn’s ‘SecondLevelModel’ function in Python (i) a one-sample t-test on all 61 subjects was used to assess whether the group-level effect significantly differs from zero (ii) an unpaired, two-sample t-test was applied to investigate the contrast difference between two groups of subjects - namely, a group of meditators (n=30) and a control group (n=31). The resulting z-statistic maps from the second-level analyses were corrected for multiple comparisons using the False Discovery Rate (FDR) method at an alpha level of 0.05, and voxels surviving FDR correction are reported and considered statistically significant in the context of our study.

#### Functional Connectivity

A priori Default Mode Network (DMN) seeds - the Posterior Cingulate Cortex (PCC) [MNI:(0,-53,26)], medial Prefrontal Cortex (mPFC) [MNI:(0,52,-6)], left Inferior Parietal Lobule (left-IPL)[MNI:(−48,-62,36)], and right Inferior Parietal Lobule (right-IPL) [MNI:(46,-62,36)] - were employed in the functional connectivity (FC) analysis^53^. The definitions of the three subnetworks of Default Network - A, B and C were extracted from the Yeo 17 Network atlas^54^ and illustrated in Figure S10. Time series from a sphere of radius 10 mm centered on these coordinates were extracted, and pairwise correlations with all other voxels in the brain were computed. For the resting state scans 4 min time series was selected (t_start_=30 s, t_end_=270 s). The 20 min long yoga nidra scan, was divided into four periods: T1 (t_start_=0s, t_end_=240s), T2 (t_start_=240s, t_end_=480s), T3 (t_start_=530s, t_end_=770s) each lasting 240 seconds, and T4(t_start_=830s, t_end_=1045s) spanning 215 seconds for connectivity analysis. These periods do not contain any explicit breathing instructions. After standardizing these time series data using their z-score, we created a correlation matrix depicting the functional connectivity (FC) between each DMN seed and all other voxels in the brain. These correlation matrices underwent Fisher-Z transformation to facilitate group-level statistical analyses. One-sample t-test was performed on all subjects (n=61) to obtain group statistics. The two-sample t-test was conducted on each matrix element to determine significant differences in functional connections between groups.

## Acknowledgements

This project was funded by Department of Science and Technology, Government of India, under the Science and Technology for Yoga and Meditation (SATYAM) scheme (sanction number DST/SATYAM/201/121(G) dated 12-09-2018).

