## Supplementary Information for "Brain Connectivity Changes in Meditators and Novices during Yoga Nidra: A Novel fMRI Study"

**This PDF file includes:**

Supplementary Figures S1 to S10
Supplementary Tables S1 to S8

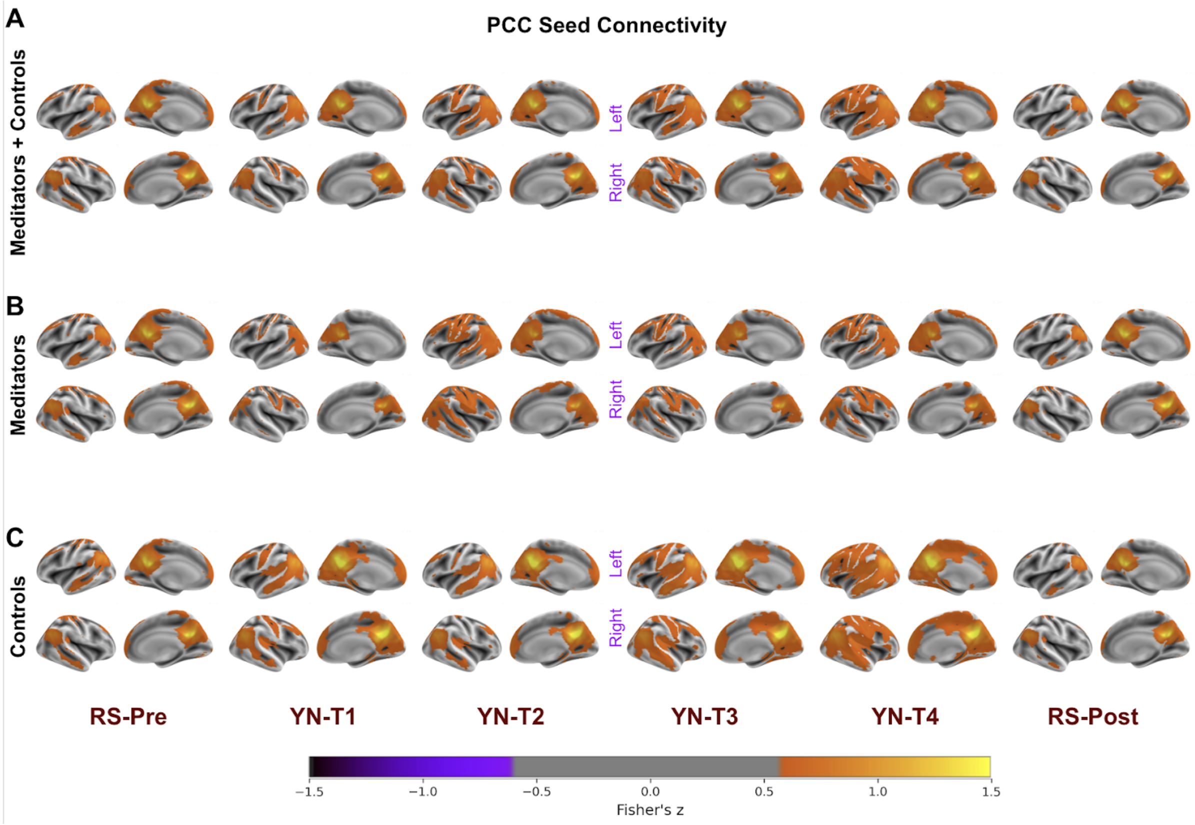

*Figure S1. Functional Connectivity (FC) maps during Resting States (RS) and Yoga Nidra Practice employing apriori DMN Node, Posterior Cingulate Cortex (PCC). The intensity of FC is displayed on surface maps using Fisher's z-value of Pearson's correlation, thresholded at z=0.6. The color scale represents the strength of the correlation, with warmer colors indicating stronger connectivity. These results demonstrate the influence of the DMN in both resting states and during Yoga Nidra.*

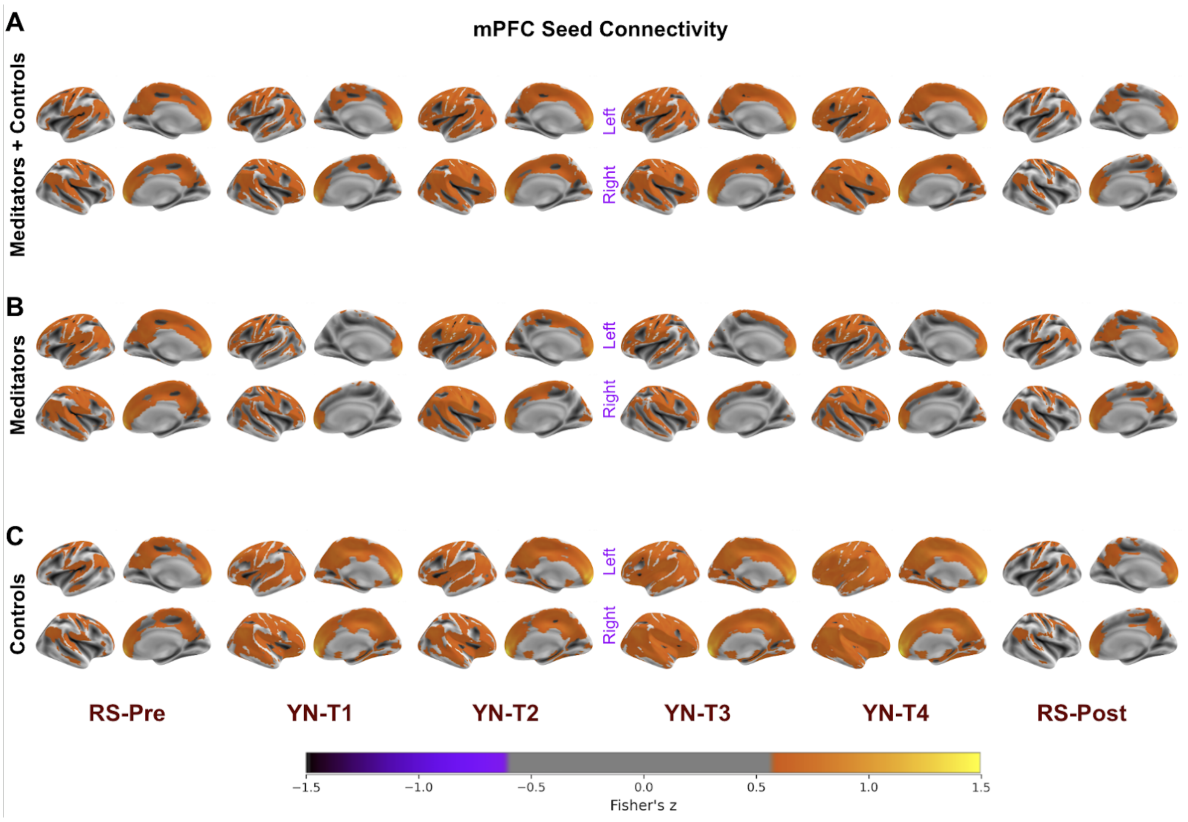

***Figure S2.*** *Functional Connectivity (FC) maps during Resting States (RS) and Yoga Nidra Practice employing medial Prefrontal Cortex (mPFC). The intensity of FC is displayed on surface maps using Fisher's z-value of Pearson's correlation, thresholded at z=0.6. The color scale represents the strength of the correlation, with warmer colors indicating stronger connectivity. These results demonstrate the influence of the DMN in both resting states and during Yoga Nidra.*

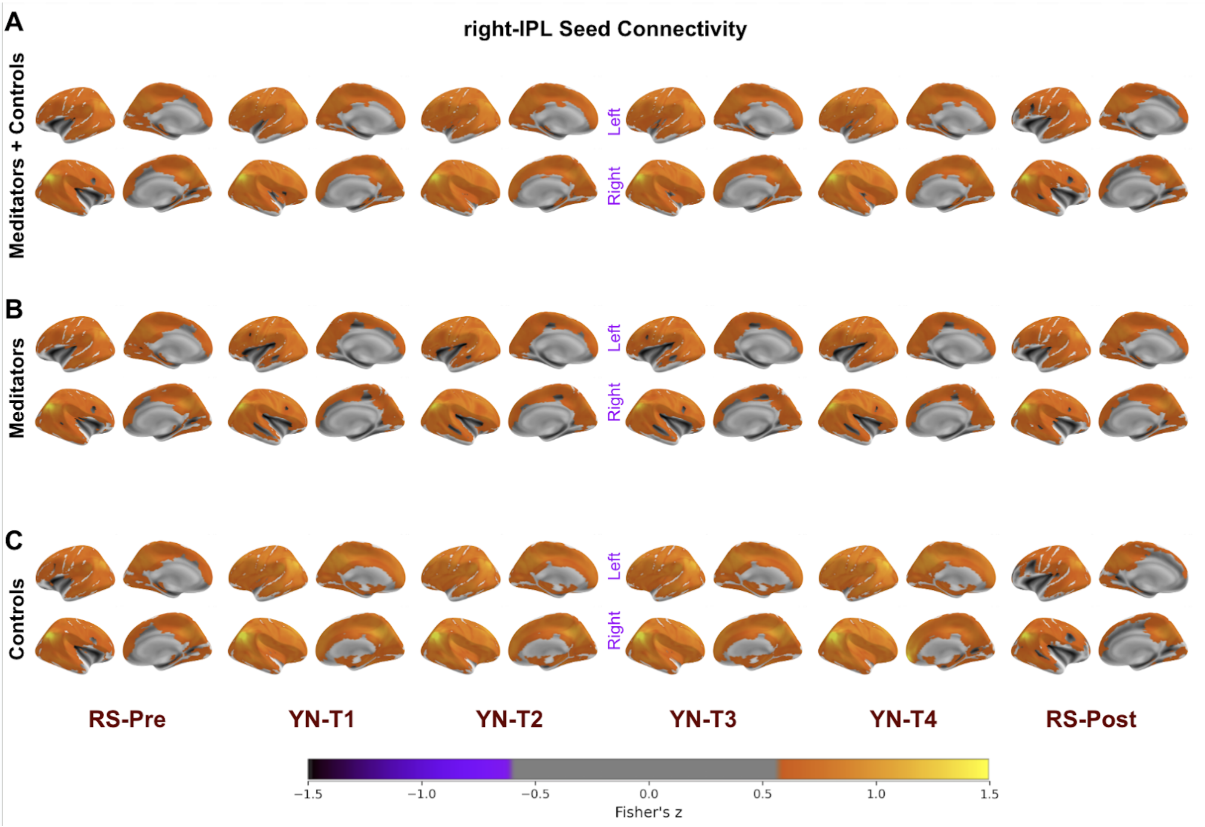

***Figure S3.*** *Functional Connectivity (FC) maps during Resting States (RS) and Yoga Nidra Practice employing right Inferior Parietal Lobule (right-IPL) [MNI:(46, -62, 36)]. The intensity of FC is displayed on surface maps using Fisher's z-value of Pearson's correlation, thresholded at z=0.6. The color scale represents the strength of the correlation, with warmer colors indicating stronger connectivity. These results demonstrate the influence of the DMN in both resting states and during Yoga Nidra.*

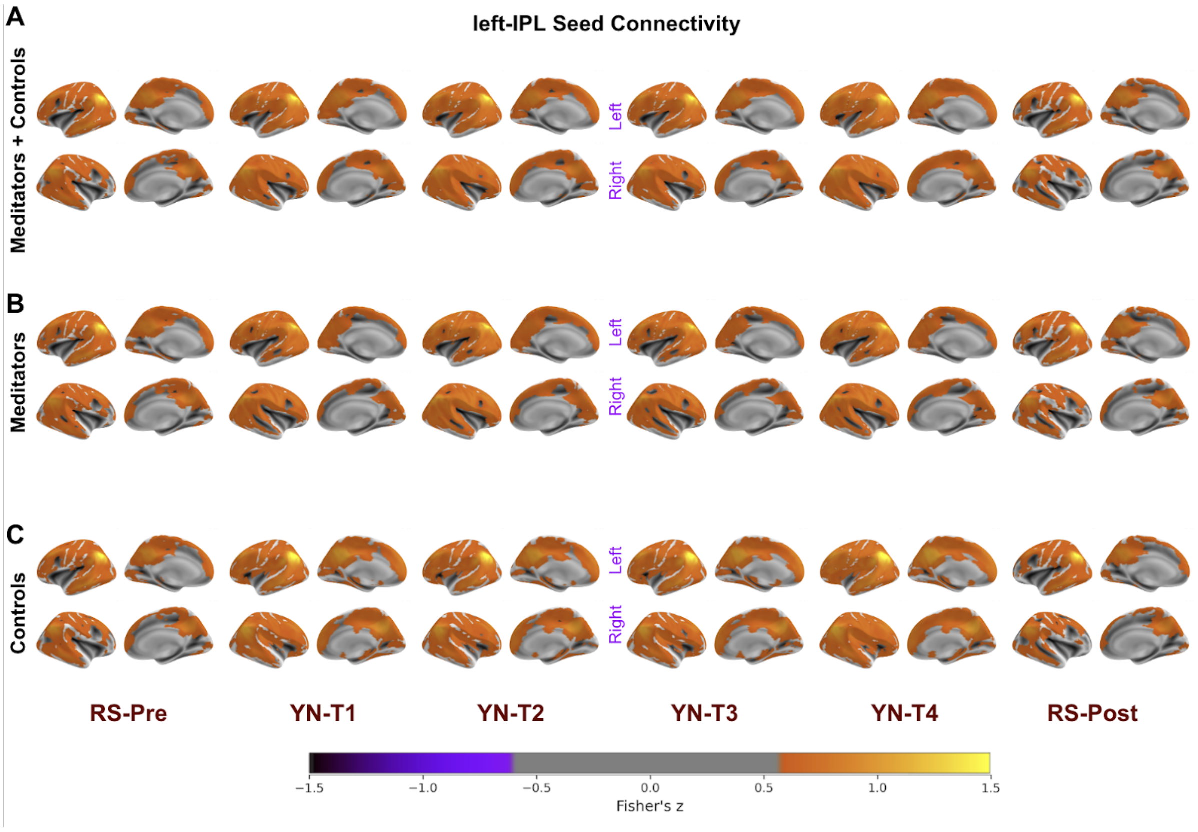

*Figure S4. Functional Connectivity (FC) maps during Resting States (RS) and Yoga Nidra Practice employing left Inferior Parietal Lobule (left-IPL)[MNI:(-48, -62, 36)]. The intensity of FC is displayed on surface maps using Fisher's z-value of Pearson's correlation, thresholded at z=0.6. The color scale represents the strength of the correlation, with warmer colors indicating stronger connectivity. These results demonstrate the influence of the DMN in both resting states and during Yoga Nidra.*

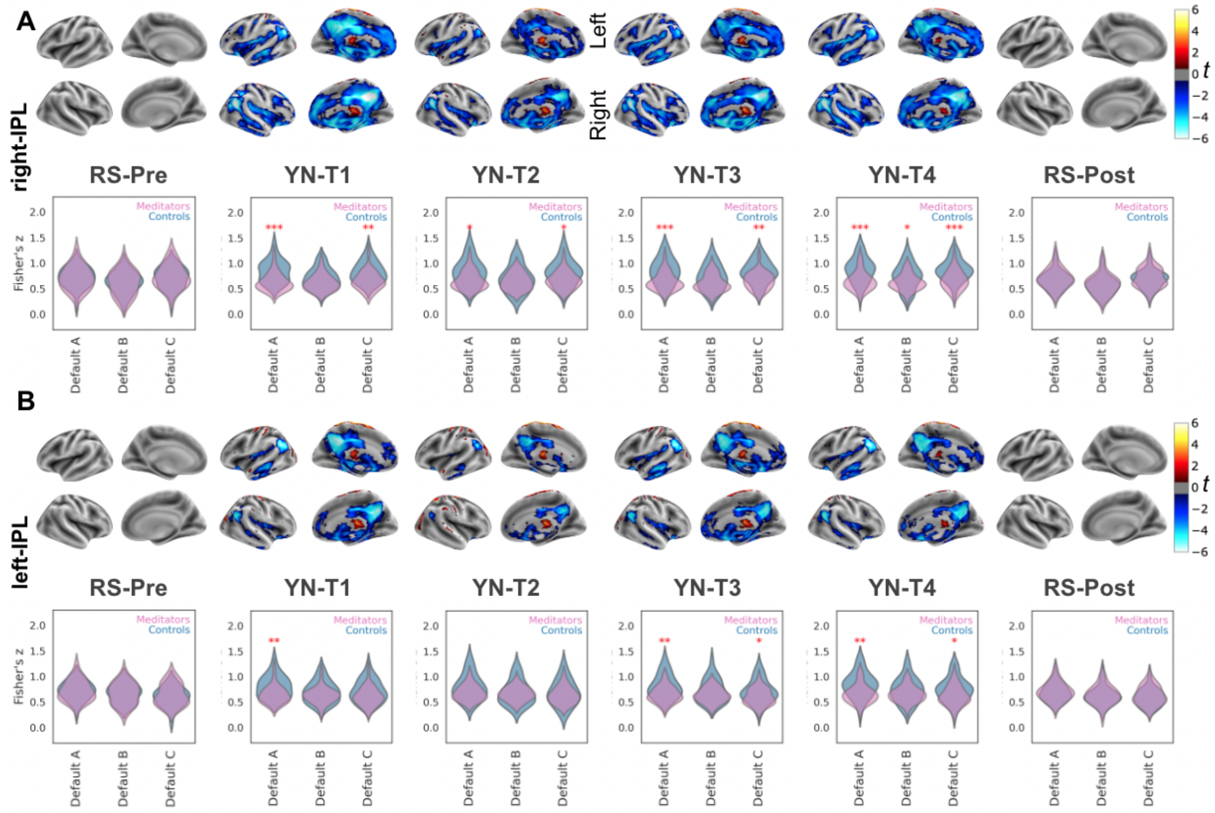

***Fig. S5.*** *Group Differences in DMN-FC between meditators and controls during Yoga Nidra but not during resting states*

*The figure uses two-sided t-tests to compare FC in meditators (n=30) and controls (n=31) using DMN seeds (A) right-IPL and (B) left-IPL. The surface plots display the t-values, corrected for multiple comparisons (FDR-corrected, q < 0.05), across the four stages of Yoga Nidra (YN), as well as in resting states pre and post-YN. The accompanying violin plots illustrate the distribution of Fisher's z-values for both groups during these stages. These plots represent the averaged Fisher's z-values of the seeds within three distinct Default Mode Network (DMN) subdivisions (Default A, Default B, and Default C), as outlined by the Schaefer Cortical Atlas. The width of the violin plot at any given y-value (Fisher's z-value) represents the proportion of data located there, providing a visual representation of the data's distribution.*

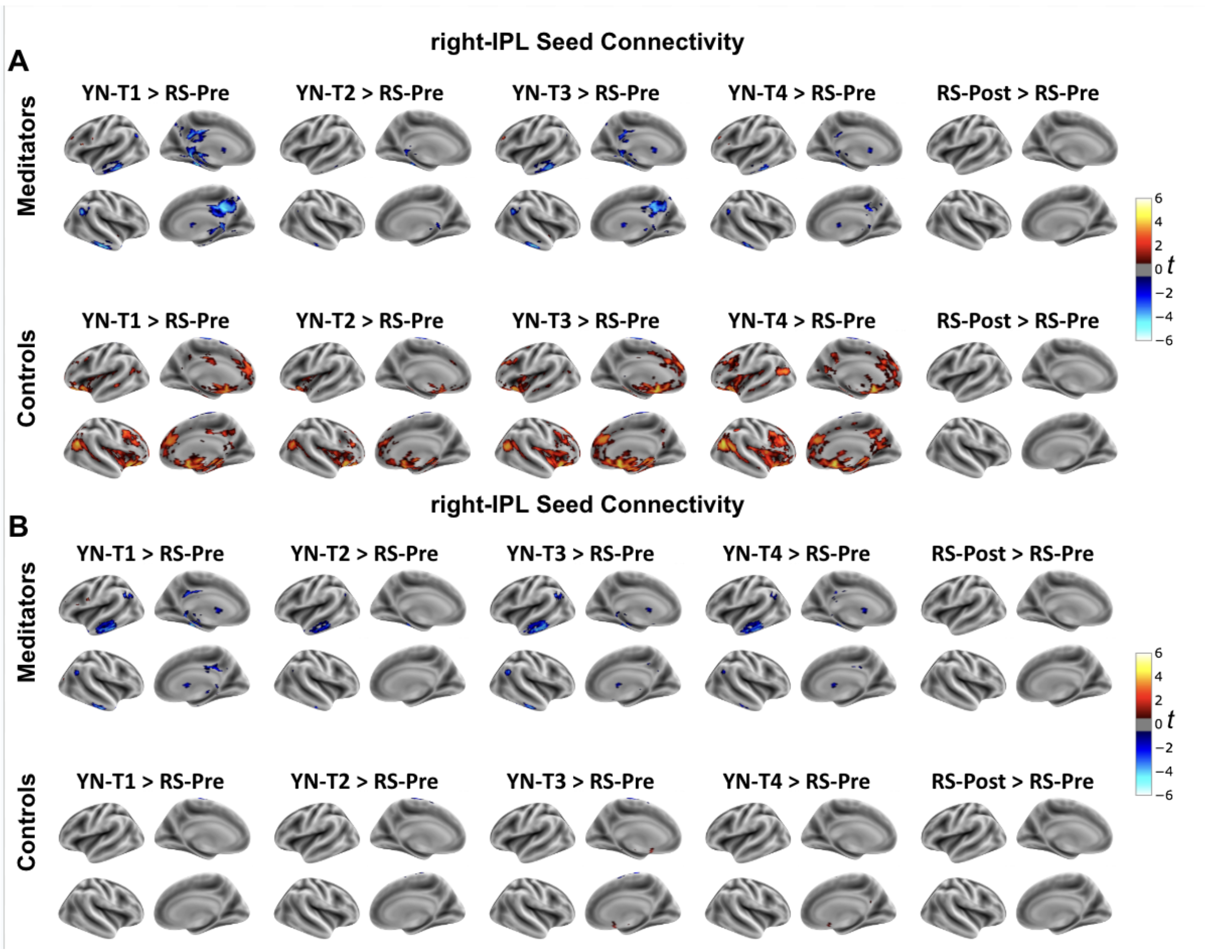

*Figure S6. Different functional connectivity changes emerge in meditators and controls during Yoga Nidra compared to rest.*

*For (A) right-IPL and (B) left-IPL seeds, a comparison of the FC during YN (T1 through T4) and the resting state post-completion of Yoga Nidra (RS-Post) is performed with resting state pre-Yoga Nidra (RS-Pre) as a baseline. The surface maps present t-values from two-sided t-tests, corrected for multiple comparisons (FDR-corrected, q < 0.05), providing a visual representation of FC differences. The figure highlights that meditators demonstrate a significant decrease in DMN connectivity during YN compared to their resting state. Conversely, controls display a significant increase in connectivity during YN relative to the resting state.*

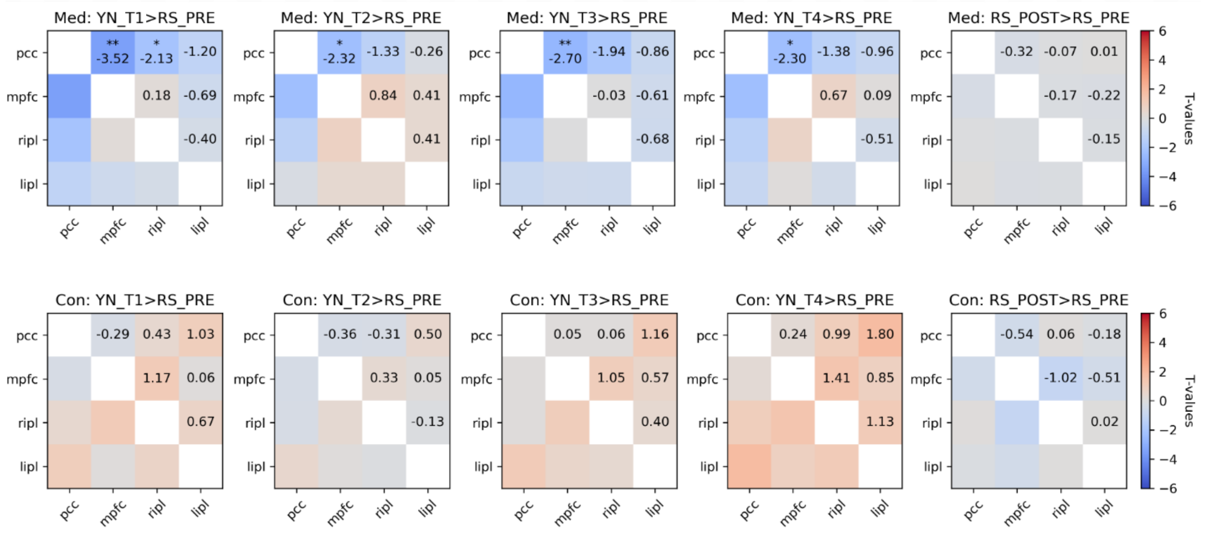

***Figure S7:*** *Intra-DMN Functional Connectivity (FC) in Meditators and Controls during Yoga Nidra Compared to Resting State.*

*The intra-DMN FC comparison involves evaluating FC during Yoga Nidra (YN) at time points T1 through T4 and the resting state post-completion of Yoga Nidra (RS-Post), using resting state pre-Yoga Nidra (RS-Pre) as the reference baseline. A t-test is conducted between the Fisher's z values of correlations among node pairs for each state and RS-Pre. The colorbars in the figure correspond to the associated t-values.*

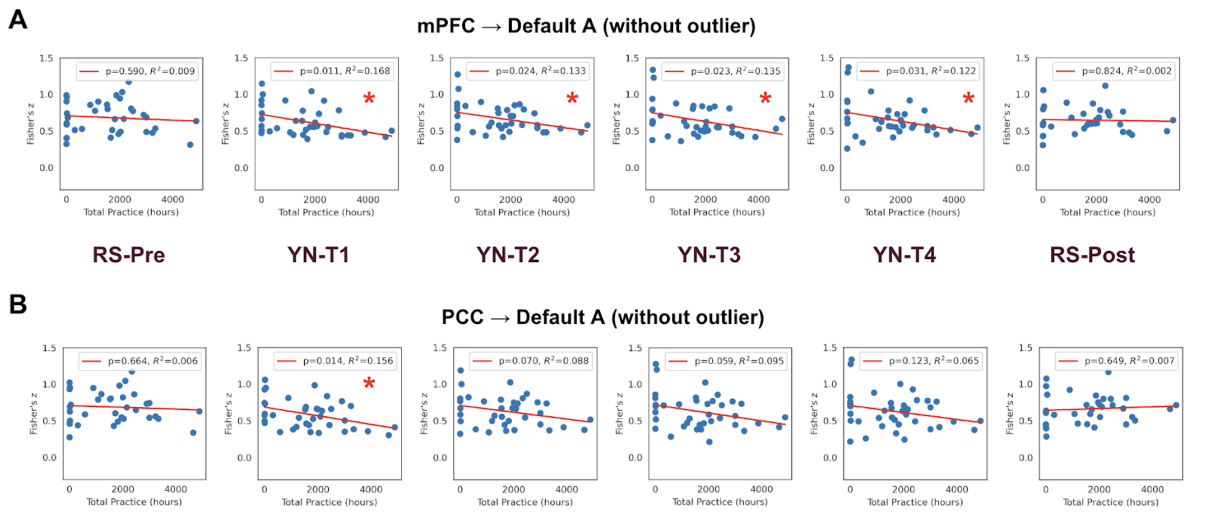

*Figure S8. Correlation between the total duration of meditation practice and the functional connectivity (FC) of (A) mPFC seed and (B) PCC seed with other regions of the DMN (Default A as defined by the Schaefer Atlas) during the Yoga Nidra (YN) practice. This analysis replicates Figure 5 from the main text, with the exception of the outlier practitioner whose cumulative practice hours exceeded 8000.*

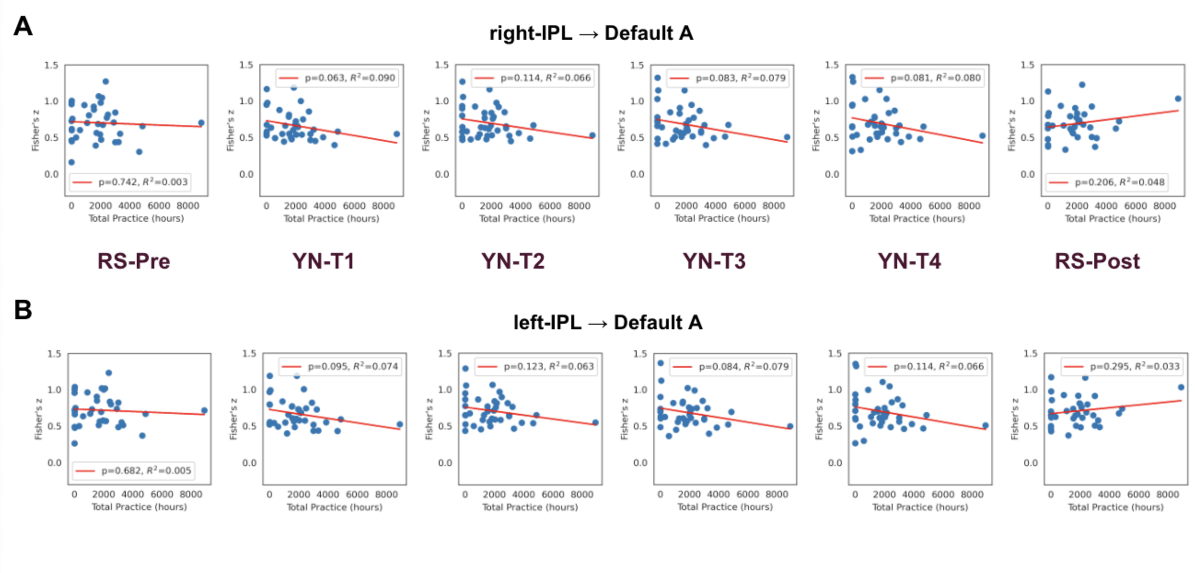

*Figure S9. Correlation between the total duration of meditation practice and the functional connectivity (FC) of (A) right-IPL seed and (B) left-IPL seed with other regions of the DMN (Default A as defined by the Schaefer Atlas) during the Yoga Nidra (YN) practice.*

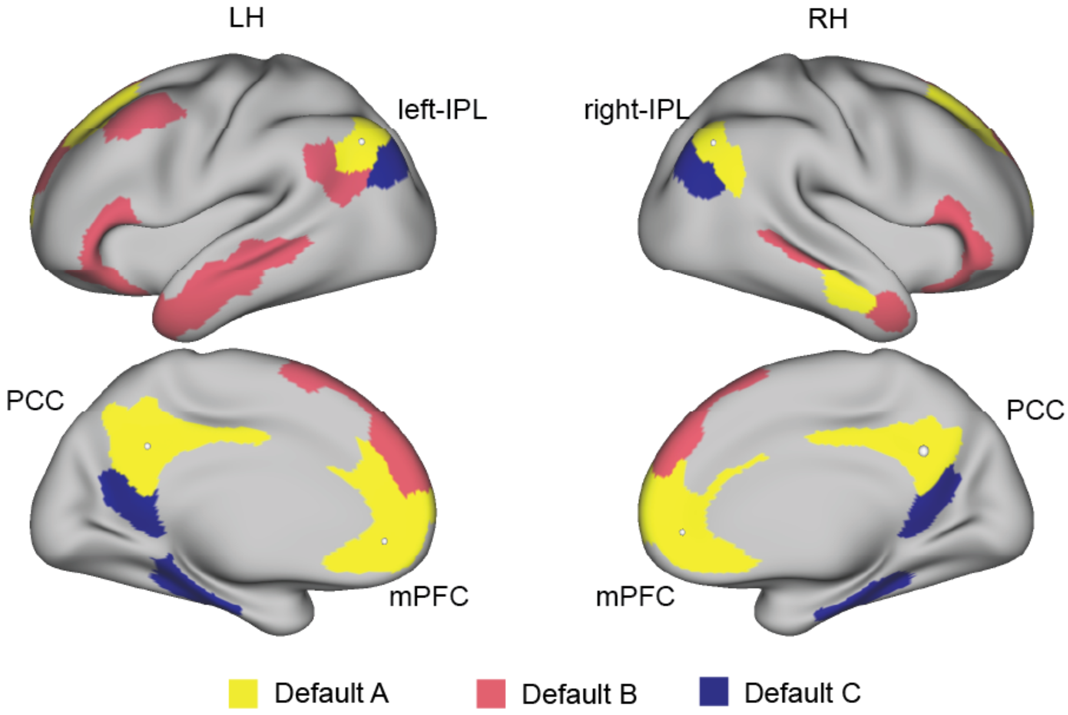

*Figure S10. Default Network and ROI definitions*

**Table S1. Demographics**

| *Demographics Variable* | Controls | Meditators |
|---|---|---|
| No. of Subjects | 31 | 30 |
| Count Female | 5 | 5 |
| Average - Age | 26.5 | 27.0 |
| Median - Family Size | 4 | 4.0 |
| Average - Body Weight (kg) | 67.0 | 67.4 |
| Average - Formal Education (years) | 17.3 | 17.0 |
| Average - Overall total practice (estimate, hrs) | 4.2 | 2,826.7 |

**Table S2. GLM analysis of Yoga Nidra, all subjects (meditators and controls collectively) revealed the following significant clusters**

| Cluster No | MNI Coordinates of peak (x, y, z) | Cluster Mean Z-score | Cluster Volume (mm3) | Regions of Activation / Deactivation |
|---|---|---|---|---|
| 1 | 60, 8, -10 | 4.39 | 1,62,496 | Auditory & Language Regions (Temporal Gyri) |
| 2 | -14, -28, -12 | 3.34 | 15,976 | Thalamus & Brain Stem |
| 3 | 4, 2, 70 | 3.29 | 13,808 | Supplementary Motor Area, Cingulate Gyrus anterior division, Postcentral Gyrus, Paracingulate Gyrus, Superior Frontal Gyrus |
| 4 | 56, 10, 42 | 3.38 | 2,784 | Precentral, Postcentral, Middle Frontal Gyrus |
| 5 | -54, 2, 48 | 3.32 | 2,720 | Precentral, Postcentral, Middle Frontal Gyrus |
| 6 | -46, -60, 26 | -3.1 | 1,784 | Middle Temporal, Angular |
| 7 | -12, 42, 54 | -3.12 | 1,608 | Superior Frontal, Frontal Sup Medial |
| 8 | 0, -78, 50 | 3.17 | 1,304 | Precuneus |
| 9 | 20, -80, 28 | -3.13 | 1,104 | Superior Occipital, Cuneus |
| 10 | 14, 46, 50 | -3.11 | 1,040 | Superior Frontal Gyrus, Frontal Pole |
| 11 | -40, -72, 52 | 3.21 | 1,000 | Angular, Parietal Sup, Sup Lateral Occipital |
| 12 | 36, -58, 24 | -2.98 | 832 | Angular |
| 13 | 10, 60, 36 | -2.98 | 800 | Superior Frontal Gyrus, Frontal Pole |
| 14 | 40, -74, 16 | -2.96 | 800 | Occipital Mid |
| 15 | -8, -52, 76 | 3.03 | 696 | Precuneus, Paracentral, Postcentral |
| 16 | -50, -38, -22 | -2.96 | 688 | Temporal Inf, Temporal Pole |
| 17 | 42, 20, -4 | 3.06 | 672 | Insula, Frontal Inf |
| 18 | -18, -60, -18 | 2.89 | 616 | Cerebellum, Brain Stem |
| 19 | 10, 56, 44 | -2.86 | 600 | Frontal Mid, Frontal Pole |
| 20 | -20, -32, -18 | -2.82 | 520 | Parahippocampal, Hippocampus |

**Table S3. PCC Seed FC difference between meditators and Controls, results of t-test within three distinct Default Mode Network (DMN) subdivisions (Default A, Default B, and Default C), as outlined by the Schaefer Cortical Atlas.**

| Stage | Network | Mean Meditators | Mean Controls | Std err Meditatots | Std err Controls | t_value | p_value |
|---|---|---|---|---|---|---|---|
| RS-Pre | DefaultA | 0.6921 | 0.6781 | 0.0396 | 0.0330 | 0.2743 | 0.7849 |
| RS-Pre | DefaultB | 0.5378 | 0.5431 | 0.0428 | 0.0353 | -0.0966 | 0.9234 |
| RS-Pre | DefaultC | 0.6643 | 0.6293 | 0.0369 | 0.0335 | 0.7048 | 0.4839 |
| YN-T1 | DefaultA | 0.5447 | 0.7656 | 0.0317 | 0.0360 | -4.6079 | 0.0000 |
| YN-T1 | DefaultB | 0.4780 | 0.5615 | 0.0302 | 0.0356 | -1.7868 | 0.0792 |
| YN-T1 | DefaultC | 0.5501 | 0.7469 | 0.0324 | 0.0386 | -3.9021 | 0.0003 |
| YN-T2 | DefaultA | 0.5982 | 0.7471 | 0.0330 | 0.0445 | -2.6902 | 0.0093 |
| YN-T2 | DefaultB | 0.5357 | 0.5560 | 0.0314 | 0.0470 | -0.3585 | 0.7213 |
| YN-T2 | DefaultC | 0.5964 | 0.7121 | 0.0341 | 0.0459 | -2.0252 | 0.0475 |
| YN-T3 | DefaultA | 0.5727 | 0.7892 | 0.0345 | 0.0415 | -4.0151 | 0.0002 |
| YN-T3 | DefaultB | 0.5020 | 0.6090 | 0.0312 | 0.0438 | -1.9910 | 0.0512 |
| YN-T3 | DefaultC | 0.5595 | 0.7435 | 0.0323 | 0.0411 | -3.5186 | 0.0009 |
| YN-T4 | DefaultA | 0.5822 | 0.8049 | 0.0346 | 0.0432 | -4.0211 | 0.0002 |
| YN-T4 | DefaultB | 0.5180 | 0.6324 | 0.0317 | 0.0450 | -2.0768 | 0.0423 |
| YN-T4 | DefaultC | 0.5860 | 0.7816 | 0.0359 | 0.0431 | -3.4857 | 0.0009 |
| RS-Pos | DefaultA | 0.6817 | 0.6444 | 0.0335 | 0.0323 | 0.7986 | 0.4280 |
| RS-Pos | DefaultB | 0.5070 | 0.5044 | 0.0381 | 0.0333 | 0.0511 | 0.9595 |
| RS-Pos | DefaultC | 0.6417 | 0.6147 | 0.0322 | 0.0305 | 0.6088 | 0.5452 |

**Table S4. mPFC Seed FC difference between meditators and Controls, results of t-test within three distinct Default Mode Network (DMN) subdivisions (Default A, Default B, and Default C), as outlined by the Schaefer Cortical Atlas.**

| Stage | Network | Mean Meditators | Mean Controls | Std err Meditatots | Std err Controls | t_value | p_value |
|---|---|---|---|---|---|---|---|
| RS-Pre | DefaultA | 0.6886 | 0.6698 | 0.0411 | 0.0338 | 0.3543 | 0.7245 |
| RS-Pre | DefaultB | 0.6065 | 0.5858 | 0.0418 | 0.0374 | 0.3703 | 0.7126 |
| RS-Pre | DefaultC | 0.5850 | 0.5612 | 0.0435 | 0.0339 | 0.4376 | 0.6634 |
| YN-T1 | DefaultA | 0.5749 | 0.7685 | 0.0307 | 0.0350 | -4.1588 | 0.0001 |
| YN-T1 | DefaultB | 0.5451 | 0.6344 | 0.0327 | 0.0339 | -1.8974 | 0.0628 |
| YN-T1 | DefaultC | 0.4874 | 0.6468 | 0.0313 | 0.0340 | -3.4525 | 0.0010 |
| YN-T2 | DefaultA | 0.6264 | 0.7476 | 0.0265 | 0.0464 | -2.2695 | 0.0270 |
| YN-T2 | DefaultB | 0.6065 | 0.6346 | 0.0281 | 0.0445 | -0.5339 | 0.5954 |
| YN-T2 | DefaultC | 0.5387 | 0.6216 | 0.0264 | 0.0436 | -1.6273 | 0.1091 |
| YN-T3 | DefaultA | 0.5897 | 0.7839 | 0.0291 | 0.0427 | -3.7552 | 0.0004 |
| YN-T3 | DefaultB | 0.5573 | 0.6753 | 0.0286 | 0.0426 | -2.2983 | 0.0252 |
| YN-T3 | DefaultC | 0.4992 | 0.6559 | 0.0276 | 0.0400 | -3.2218 | 0.0021 |
| YN-T4 | DefaultA | 0.5974 | 0.8162 | 0.0273 | 0.0432 | -4.2861 | 0.0001 |
| YN-T4 | DefaultB | 0.5708 | 0.7108 | 0.0273 | 0.0453 | -2.6468 | 0.0104 |
| YN-T4 | DefaultC | 0.5179 | 0.6810 | 0.0288 | 0.0420 | -3.1997 | 0.0022 |
| RS-Pos | DefaultA | 0.6657 | 0.6366 | 0.0313 | 0.0301 | 0.6680 | 0.5070 |
| RS-Pos | DefaultB | 0.5619 | 0.5459 | 0.0346 | 0.0342 | 0.3272 | 0.7448 |
| RS-Pos | DefaultC | 0.5594 | 0.5231 | 0.0334 | 0.0276 | 0.8466 | 0.4009 |

**Table S5. right-IPL Seed FC difference between meditators and Controls, results of t-test within three distinct Default Mode Network (DMN) subdivisions (Default A, Default B, and Default C), as outlined by the Schaefer Cortical Atlas**

| Stage | Network | Mean Meditators | Mean Controls | Std err Meditatots | Std err Controls | t_value | p_value |
|---|---|---|---|---|---|---|---|
| RS-Pre | DefaultA | 0.7164 | 0.7221 | 0.0447 | 0.0365 | -0.0998 | 0.9208 |
| RS-Pre | DefaultB | 0.6158 | 0.6248 | 0.0469 | 0.0369 | -0.1518 | 0.8799 |
| RS-Pre | DefaultC | 0.7380 | 0.7159 | 0.0457 | 0.0382 | 0.3742 | 0.7097 |
| YN-T1 | DefaultA | 0.6383 | 0.8463 | 0.0322 | 0.0400 | -4.0481 | 0.0002 |
| YN-T1 | DefaultB | 0.6181 | 0.7032 | 0.0289 | 0.0368 | -1.8210 | 0.0738 |
| YN-T1 | DefaultC | 0.6620 | 0.8348 | 0.0326 | 0.0379 | -3.4555 | 0.0010 |
| YN-T2 | DefaultA | 0.6728 | 0.8236 | 0.0319 | 0.0476 | -2.6296 | 0.0109 |
| YN-T2 | DefaultB | 0.6447 | 0.6944 | 0.0292 | 0.0465 | -0.9055 | 0.3689 |
| YN-T2 | DefaultC | 0.6749 | 0.8168 | 0.0308 | 0.0453 | -2.5887 | 0.0122 |
| YN-T3 | DefaultA | 0.6474 | 0.8464 | 0.0314 | 0.0458 | -3.5809 | 0.0007 |
| YN-T3 | DefaultB | 0.6143 | 0.7184 | 0.0301 | 0.0445 | -1.9378 | 0.0575 |
| YN-T3 | DefaultC | 0.6640 | 0.8326 | 0.0307 | 0.0404 | -3.3192 | 0.0016 |
| YN-T4 | DefaultA | 0.6576 | 0.8713 | 0.0338 | 0.0448 | -3.8078 | 0.0003 |
| YN-T4 | DefaultB | 0.6285 | 0.7380 | 0.0287 | 0.0453 | -2.0403 | 0.0459 |
| YN-T4 | DefaultC | 0.6783 | 0.8638 | 0.0346 | 0.0387 | -3.5770 | 0.0007 |
| RS-Pos | DefaultA | 0.7055 | 0.6787 | 0.0390 | 0.0329 | 0.5298 | 0.5984 |
| RS-Pos | DefaultB | 0.5940 | 0.5753 | 0.0402 | 0.0339 | 0.3592 | 0.7208 |
| RS-Pos | DefaultC | 0.7222 | 0.6883 | 0.0386 | 0.0319 | 0.6818 | 0.4983 |

**Table S6. left-IPL Seed FC difference between meditators and Controls, results of t-test within three distinct Default Mode Network (DMN) subdivisions (Default A, Default B, and Default C), as outlined by the Schaefer Cortical Atlas**

| Stage | Network | Mean Meditators | Mean Controls | Std err Meditatots | Std err Controls | t_value | p_value |
|---|---|---|---|---|---|---|---|
| RS-Pre | DefaultA | 0.7257 | 0.7247 | 0.0396 | 0.0336 | 0.0184 | 0.9854 |
| RS-Pre | DefaultB | 0.6633 | 0.6688 | 0.0416 | 0.0337 | -0.1035 | 0.9179 |
| RS-Pre | DefaultC | 0.6462 | 0.6023 | 0.0408 | 0.0351 | 0.8197 | 0.4159 |
| YN-T1 | DefaultA | 0.6439 | 0.8045 | 0.0317 | 0.0399 | -3.1479 | 0.0026 |
| YN-T1 | DefaultB | 0.6311 | 0.6947 | 0.0285 | 0.0386 | -1.3262 | 0.1900 |
| YN-T1 | DefaultC | 0.6108 | 0.7109 | 0.0321 | 0.0403 | -1.9446 | 0.0567 |
| YN-T2 | DefaultA | 0.6832 | 0.7896 | 0.0276 | 0.0462 | -1.9762 | 0.0529 |
| YN-T2 | DefaultB | 0.6657 | 0.6872 | 0.0247 | 0.0438 | -0.4269 | 0.6711 |
| YN-T2 | DefaultC | 0.6352 | 0.6835 | 0.0297 | 0.0470 | -0.8685 | 0.3887 |
| YN-T3 | DefaultA | 0.6527 | 0.8124 | 0.0290 | 0.0425 | -3.1046 | 0.0029 |
| YN-T3 | DefaultB | 0.6375 | 0.7145 | 0.0263 | 0.0425 | -1.5382 | 0.1294 |
| YN-T3 | DefaultC | 0.6059 | 0.7093 | 0.0294 | 0.0415 | -2.0321 | 0.0467 |
| YN-T4 | DefaultA | 0.6616 | 0.8383 | 0.0327 | 0.0456 | -3.1518 | 0.0026 |
| YN-T4 | DefaultB | 0.6555 | 0.7317 | 0.0282 | 0.0446 | -1.4438 | 0.1542 |
| YN-T4 | DefaultC | 0.6249 | 0.7435 | 0.0345 | 0.0436 | -2.1319 | 0.0373 |
| RS-Pos | DefaultA | 0.7101 | 0.6978 | 0.0356 | 0.0322 | 0.2551 | 0.7996 |
| RS-Pos | DefaultB | 0.6393 | 0.6291 | 0.0364 | 0.0330 | 0.2087 | 0.8355 |
| RS-Pos | DefaultC | 0.6306 | 0.5934 | 0.0359 | 0.0315 | 0.7815 | 0.4379 |

**Table S7: 2-way ANOVA Post hoc differences table**

| | coef | std err | t | P>\|t\| | 0.025 | 0.975 |
|---|---|---|---|---|---|---|
| **Intercept** | 0.6366 | 0.035 | 18.163 | 0 | 0.568 | 0.706 |
| **C(Group)[T.Meditators]** | 0.0291 | 0.051 | 0.565 | 0.572 | -0.072 | 0.13 |
| **C(Stage)[T.RS_PRE]** | 0.0332 | 0.05 | 0.67 | 0.503 | -0.064 | 0.131 |
| **C(Stage)[T.YN_T1]** | 0.1319 | 0.05 | 2.661 | 0.008 | 0.034 | 0.229 |
| **C(Stage)[T.YN_T2]** | 0.111 | 0.05 | 2.239 | 0.026 | 0.013 | 0.208 |
| **C(Stage)[T.YN_T3]** | 0.1473 | 0.05 | 2.971 | 0.003 | 0.05 | 0.245 |
| **C(Stage)[T.YN_T4]** | 0.1796 | 0.05 | 3.624 | 0 | 0.082 | 0.277 |
| **C(Group)[T.Meditators]:C(Stage)[T.RS_PRE]** | -0.0104 | 0.072 | -0.143 | 0.886 | -0.153 | 0.132 |
| **C(Group)[T.Meditators]:C(Stage)[T.YN_T1]** | -0.2227 | 0.071 | -3.117 | 0.002 | -0.363 | -0.082 |
| **C(Group)[T.Meditators]:C(Stage)[T.YN_T2]** | -0.1503 | 0.071 | -2.104 | 0.036 | -0.291 | -0.01 |
| **C(Group)[T.Meditators]:C(Stage)[T.YN_T3]** | -0.2232 | 0.071 | -3.125 | 0.002 | -0.364 | -0.083 |
| **C(Group)[T.Meditators]:C(Stage)[T.YN_T4]** | -0.2479 | 0.071 | -3.471 | 0.001 | -0.388 | -0.107 |
| Omnibus: | 10.683 | Durbin-Watson: | 2.008 | | | |
| Prob(Omnibus): | 0.005 | Jarque-Bera (JB): | 10.859 | | | |
| Skew: | 0.424 | Prob(JB): | 0.00439 | | | |
| Kurtosis: | 3.144 | Cond. No. | 18.1 | | | |

**Table S8: Multi-band factors across the resting state and Yoga Nidra protocols across subjects.**

| Subject Code | Group | Resting State | Yoga Nidra | | Subject Code | Group | Resting State | Yoga Nidra |
|---|---|---|---|---|---|---|---|---|
| sub-00 | Meditators | 4 | 8 | | sub-c01 | Controls | 4 | 3 |
| sub-01 | Meditators | 4 | 8 | | sub-c02 | Controls | 4 | 3 |
| sub-02 | Meditators | 4 | 8 | | sub-c03 | Controls | 4 | 3 |
| sub-03 | Meditators | 4 | 8 | | sub-c04 | Controls | 4 | 3 |
| sub-04 | Meditators | 4 | 8 | | sub-c05 | Controls | 4 | 3 |
| sub-05 | Meditators | 4 | 8 | | sub-c06 | Controls | 4 | 3 |
| sub-06 | Meditators | 4 | 8 | | sub-c07 | Controls | 4 | 3 |
| sub-07 | Meditators | 4 | 8 | | sub-c08 | Controls | 4 | 3 |
| sub-08 | Meditators | 4 | 8 | | sub-c09 | Controls | 4 | 3 |
| sub-09 | Meditators | 4 | 8 | | sub-c10 | Controls | 4 | 3 |
| sub-10 | Meditators | 4 | 8 | | sub-c11 | Controls | 4 | 3 |
| sub-11 | Meditators | 4 | 8 | | sub-c12 | Controls | 4 | 3 |
| sub-12 | Meditators | 4 | 8 | | sub-c13 | Controls | 4 | 3 |
| sub-13 | Meditators | 4 | 8 | | sub-c14 | Controls | 4 | 3 |
| sub-14 | Meditators | 4 | 8 | | sub-c15 | Controls | 4 | 3 |
| sub-15 | Meditators | 4 | 8 | | sub-c16 | Controls | 4 | 3 |
| sub-16 | Meditators | 4 | 8 | | sub-c17 | Controls | 4 | 3 |
| sub-17 | Meditators | 4 | 8 | | sub-c18 | Controls | 4 | 3 |
| sub-18 | Meditators | 4 | 8 | | sub-c19 | Controls | 4 | 3 |
| sub-19 | Meditators | 4 | 8 | | sub-c20 | Controls | 4 | 3 |
| sub-20 | Meditators | 4 | 8 | | sub-c21 | Controls | 4 | 3 |
| sub-21 | Meditators | 4 | 8 | | sub-c22 | Controls | 4 | 3 |
| sub-22 | Meditators | 4 | 8 | | sub-c23 | Controls | 4 | 3 |
| sub-23 | Meditators | 4 | 8 | | sub-c24 | Controls | 4 | 3 |
| sub-24 | Meditators | 4 | 8 | | sub-c25 | Controls | 4 | 3 |
| sub-25 | Meditators | 4 | 8 | | sub-c26 | Controls | 4 | 3 |
| sub-26 | Meditators | 4 | 8 | | sub-c27 | Controls | 4 | 3 |
| sub-27 | Meditators | 4 | 3 | | sub-c28 | Controls | 4 | 3 |
| sub-28 | Meditators | 4 | 3 | | sub-c29 | Controls | 4 | 3 |
| sub-29 | Meditators | 4 | 3 | | sub-c30 | Controls | 4 | 3 |
| | | | | | sub-c31 | Controls | 4 | 3 |
